# Reporter-based screening identifies RAS-RAF mutations as drivers of resistance to active-state RAS inhibition in colorectal cancer

**DOI:** 10.1101/2024.07.01.601542

**Authors:** Oleksandra Aust, Eric Blanc, Mareen Lüthen, Viola Hollek, Rosario Astaburuaga-García, Bertram Klinger, Alexandra Trinks, Dieter Beule, Björn Papke, David Horst, Nils Blüthgen, Christine Sers, Channing J. Der, Markus Morkel

**Author notes:** Corresponding author: Markus Morkel.

## Abstract

Therapy-induced acquired resistance limits the clinical effectiveness of mutation-specific RAS inhibitors in colorectal cancer. It is unknown whether broad-spectrum active-state RAS inhibitors meet similar limitations. Here, we identify and categorize mechanisms of resistance to the broad-spectrum active-state RAS inhibitor RMC-7977 in colorectal cancer cell lines. We found that KRAS-mutant colorectal cancer cell lines are universally sensitive to RMC-7977, inhibiting the RAS-RAF-MEK-ERK axis, halting proliferation and in some cases inducing apoptosis. To monitor KRAS downstream effector pathway activity, we developed a compartment-specific dual-color ERK activity reporter. RMC-7977 treatment reduced reporter activity. However, long-term dose escalation with RMC-7977 revealed multiple patterns of reporter reactivation in emerging resistant cell populations that correlated with phosphorylation states of compartment-specific ERK targets. Cells sorted for high, low, or cytoplasmic reporter activity exhibited distinct patterns of genomic mutations, phospho-protein, and transcriptional activities. Notably, all resistant subpopulations showed dynamic ERK regulation in the presence of the RAS inhibitor, unlike the parental sensitive cell lines. High levels of RAS downstream activities were observed in cells characterized by a KRAS Y71H resistance mutation. In contrast, RAS inhibitor-resistant populations with low, or cytoplasmic ERK reporter reactivation displayed different genetic alterations, among them RAF1 S257L and S259P mutations. Colorectal cancer cells resistant to RMC-7977 and harboring the RAF1 mutation specifically exhibited synergistic sensitivity to concurrent RAS and RAF inhibition. Our findings endorse reporter-assisted screening together with single-cell analyses as a powerful approach for dissecting the complex landscape of therapy resistance. The strategy offers opportunities to develop clinically relevant combinatorial treatments to counteract emergence of resistant cancer cells.

## Introduction

RAS proto-oncogenes encode small GTPases that relay signals from receptors such as receptor tyrosine kinases to multiple downstream pathways (*1*). Oncogenic mutant forms of RAS are among the most common drivers of human cancer. The mutations, which mostly occur in the codon 12, 13 and 61 hotspots of KRAS, NRAS, or HRAS stabilize the active GTP-bound state, and thereby increase engagement of RAS downstream pathways that endow cells with cancer hallmarks such as self-sufficient proliferation and resistance to apoptosis (*2*). A central pathway activated by RAS is the mitogen-activated protein kinase (MAPK) cascade with ERK1 and ERK2 (together abbreviated as ERK hereafter) as terminal kinases that activate distinct sets of cytoplasmic and nuclear targets (*3, 4*).

Approximately 40% of colorectal cancers (CRCs) harbour an oncogenic KRAS hotspot mutation, while another 5% hold NRAS mutations (*5*). Currently, patients without mutations in the MAPK cascade or with BRAF mutations profit from therapies targeting EGFR or EGFR/RAF, respectively (*6*), but patients with RAS mutations are excluded from targeted therapy. This is because targeted inhibition of MAPK in RAS-activated CRC usually resulted in the re-activation of oncogenic signals due to feedback mechanisms (*7, 8*). Therefore, it has been suggested that direct targeting of RAS proteins would be most effective, but RAS-family GTPases have long withstood selective inhibition by small molecules (*9*). Only recently, KRAS inhibitors have become available that exclusively target inactive-state mutant KRAS G12C and G12D protein (*10–12*). Clinical impact of these inhibitors was limited in CRC, as patients harbouring the specific mutations are rare, and, furthermore, patients rapidly developed clinical resistance (*13, 14*).

Explorations into secondary resistance against mutation-specific inactive-state KRAS G12C/D inhibitors uncovered multiple mechanisms. These encompassed the acquisition and clonal expansion of new oncogenic KRAS mutations, as found in clinical studies (*14, 15*) as well as in *in-vitro* experiments (*14–16*), mutations in other components of the MAPK pathway (*14*), for instance in receptor tyrosine kinases such as EGFR or MET or in RAS downstream effectors, such as MAP2K1 encoding MEK1. Additionally, resistance drivers that act independently or in alongside to RAS downstream pathways were detected, such as histological transformation or activation of YAP-TEAD signalling (*17, 18*). The former mechanisms re-activate the bottom-most MAPK kinase ERK, whereas the latter mechanisms are thought to occur in the absence of detectable ERK reactivation.

Beyond targeting inactive-state RAS, a more recent strategy is to target the active state of oncogenic RAS by the non-covalent assembly of a trimeric complex, forming between an inhibitor that first binds to the ubiquitous and essential protein cyclophilin A and ultimately complexes with active-state RAS (*19*). An optimized structure-guided design yielded the first broad-spectrum active-state RAS inhibitor RMC-7977 (*20*). Initial studies have shown broad anti-cancer activity of RMC-7797 and a favourable toxicity profile in RAS-addicted models of pancreatic cancer (*20, 21*). However, sensitivity to RMC-7977 in KRAS-mutant CRC is unknown and potential secondary resistance mechanisms have not been identified.

Here, we employed KRAS-mutant CRC cell lines to assess sensitivities and unravel resistance mechanisms to RMC-7977. Using a reporter-based screening approach, we found that the emergence of resistance to broad-spectrum active-state RAS inhibition in CRC cell lines can be due to functionally different mechanisms resulting in different strengths of MAPK-and PI3K-downstream pathway reactivation. We identified mutations driving resistance to active-state RAS inhibition centring on KRAS and RAF1 and overlapping considerably with pathogenic gene variants causing RASopathies, thus clearly deviating from mutations driving resistance to inactive-state RAS inhibitors.

## Results

### RAS-mutant CRC cell lines are universally sensitive to active-state RAS inhibition

To examine the effect of broad-spectrum RAS inhibition in CRC, we employed a panel of CRC cell lines. We found that nine CRC cell lines carrying hotspot mutations in KRAS were all growth-inhibited by the active-state RAS inhibitor RMC-7977 (RASi) in the nanomolar range (Fig. 1A), while two CRC cell lines bearing BRAF^V600E^ mutations were insensitive to RASi. The ERK inhibitor SCH772984 (*22*) (ERKi) likewise inhibited growth of KRAS-mutant cell lines, albeit at higher concentrations, and LS174T and SW837 cells, bearing G12D and G12C mutations, respectively, were in addition sensitive to mutation-specific off-state RAS inhibitors (Fig. 1B). To test whether RMC-7977 was also cytotoxic, we tested for apoptosis. In SW480, LS174T and SW837 cells, but not in DLD1, growth inhibition by RMC-7977 was accompanied by cell death, as assessed by quantification of cleaved Caspase-3-positive cells (Fig. 1C). In all KRAS-mutant CRC cell lines tested, RMC-7977 inhibited phosphorylation of the terminal MAPK kinase ERK at doses as low as 1 to 10 nM (Fig. 1D). The data show that broad-spectrum active-state RAS inhibition results in universal blockade of MAPK signalling and cancer cell proliferation in KRAS-mutant CRC cell lines.

**Figure 1:**
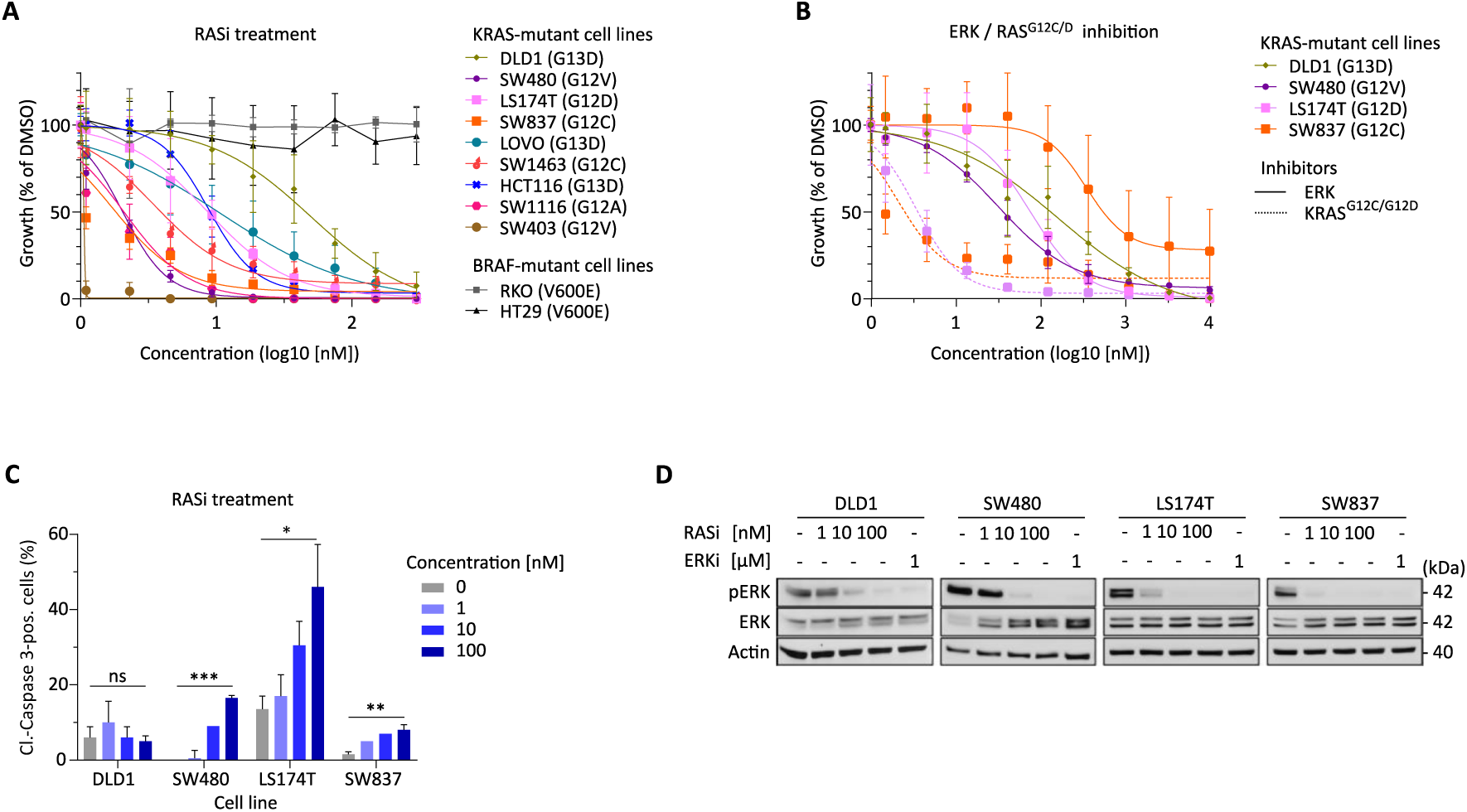
RMC-7977 blocks proliferation and ERK phosphorylation in KRAS-mutant CRC cell lines. **A-B** Proliferation (nuclear count) of cell lines upon drug treatment, as indicated, mean of n=3 replicate experiments. **A** Response to RMC-7977 (RASi) in CRC cell lines, KRAS- and BRAF-mutations as indicated. **B** Response to site-specific RAS inhibitors MRTX849 and MRTX1133 or ERK inhibitor SCH772984 (ERKi), as indicated. **C** Induction of apoptosis in response to RASi in CRC cell lines, as measured by positivity for cleaved Caspase-3, n=2 replicate experiments, ANOVA significance tests. ns, not significant; *, p<0.05; **, p<0.01; ***, p<0.001 **D** ERK phosphorylation in response to 1 h treatment with RAS (RMC-7797) or ERK (SCH772984) inhibitor in CRC cell lines.

### An optimized degradation-based reporter of cytoplasmic and nuclear ERK activity

To follow the dynamics of MAPK pathway activity during RAS inhibitor treatment and acquisition of secondary resistance, we employed fluorescent ERK activity monitoring. Our aim was to measure integrated ERK activity over several-hour intervals rather than short-term fluctuations, and additionally, we wanted to distinguish between cytoplasmic and nuclear ERK activity. To achieve this, we developed a pair of novel ERK reporters, which we termed FIREX, as they represent an extension of the published fluorescent integrative reporter of ERK, FIRE (*23*).

For the construction of these reporters, we used the high-activity EF-1a promoter to drive transgenes composed of the Myc nuclear localisation signal or the IkBa nuclear exclusion signal, fused to red fluorescent mKate2 or green fluorescent mVenus proteins, respectively, along with the Fra1 degron (Fig. 2A). This design ensured regulation of fluorescent proteins in an ERK activity-dependent manner, as in the original FIRE reporter. Upon transfection into HEK 293 cells, the constructs elicited localized nuclear red or cytoplasmic green fluorescence signals, indicating functionality of the localisation signals (Fig 2B). The red nuclear and green cytoplasmic reporters were termed nFIREX and cFIREX, respectively.

**Figure 2:**
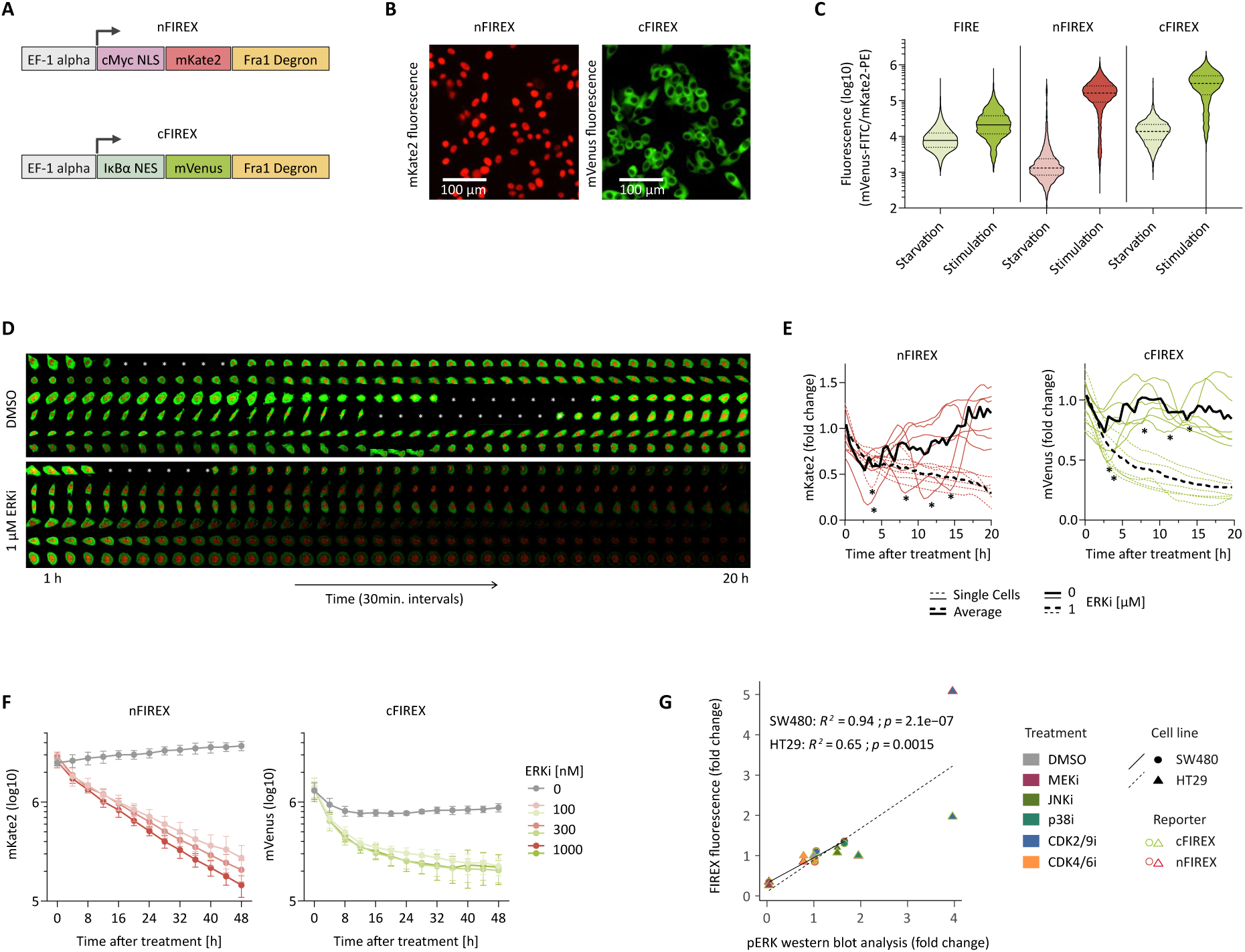
An optimized degradation-based reporter of cytoplasmic and nuclear ERK activity. **A** Domain structure of the nFIREX and cFIREX reporters. The FRA-1 degron contains ERK phosphorylation sites, as also seen in Fig. 4A. **B** Fluorescent signals elicited by FIRE, nFIREX and cFIREX after transfection into HEK293ΔRAF1:ER cells. **C** Dynamic range of FIRE(*24*), nFIREX and cFIREX in tamoxifen-inducible HEK293ΔRAF1:ER cells. Reporter signals are measured by flow cytometry 24 h after FCS starvation and tamoxifen stimulation. **D** Time-lapse microscopy of SW480 FIREX cells, ± addition of 1 μM ERKi at timepoint of first images taken. **E** Quantification of FIREX signals from images as in (D) on single-cell basis, using Cell Pathfinder. * indicate mitoses. **F** Quantification of FIREX response to increasing doses of ERKi in SW480 cells on a population basis using Incucyte. **G** Specificity tests of FIREX in SW480 and HT29 cells, inhibitors as indicated. Correlation plot shows quantification of FIREX response 16h after inhibitor treatment in stably FIREX-transfected SW480 and HT29 cells, using Incucyte and quantification of phospho-ERK by western blot analysis, 1h after inhibitor treatment. Pearson correlation coefficients and p-values are given per cell line.

To evaluate the responsiveness of the reporters to changes in MAPK activity, we stably transfected them into HEK293 RAF:ER cells(*25*), which can be induced for active RAF1 by tamoxifen. We then measured reporter fluorescence after serum starvation and subsequent induction of constitutively active RAF1 for 24 h. While the activity of the original FIRE reporter increased 7-fold after induction, fluorescence elicited by the nFIREX and cFIREX reporters increased 38-fold and 20-fold, respectively (Fig 2C). These data show that FIREX responds to the induction of ERK activity similar to FIRE, but with a greater dynamic range.

We followed reporter activities of FIREX in stably transfected SW480 CRC cells, and in the presence or absence of ERKi (Fig. 2D-F). As expected, activities in individual cells fluctuated during the cell cycle, and were strongly repressed by the ERK inhibitor (Fig. 2D, quantified in 2E). On a population level, nFIREX activity showed a nearly linear decrease for 48 h after ERK inhibitor treatment, while cFIREX activity reached a plateau after 24 h, likely due to higher autofluorescence in the green channel that capped the lower limit of the dynamic range (Fig. 2F).

To assess the specificity of the FIREX reporters, we compared reporter activities with Western blot analyses obtained after treatment with inhibitors of MEK, JNK, p38, CDK2/9 and CDK4/6, and found significant correlations (Fig. 2G; Supplementary Fig. 1). In both SW480 and HT29 CRC cells, the MEK inhibitor decreased both FIREX activities and ERK phosphorylation. Unexpectedly, the treatment with CDK2/9 inhibitor resulted in increased FIREX activities in HT29 cells, a pattern mirrored by increased ERK phosphorylation in the Western blot. Most other treatments induced only minor changes in reporter activity and ERK phosphorylation, underscoring the specificity of the FIREX reporter system for ERK1/2.

### Tracking of ERK activity during development of RAS inhibitor resistance in CRC cells

To monitor ERK activity during the emergence of resistance to active-state RAS inhibition, we stably transfected SW480 and DLD1 cells with the FIREX plasmids. We established monoclonal populations of both cell lines by sorting single cells by fluorescence, expansion, and selection of clonal populations with dynamic reporter activity. Subsequently, we exposed the cells to RASi concentrations that partially inhibited growth (1 nM for SW480 and 20 nM for DLD1, corresponding to the IC30 values for the cell lines). Over a period of 2 to 4 months, we performed dose-escalation experiments to generate SW480 and DLD1 cell populations capable of tolerating 16 nM and 160 nM RMC-7977, respectively. Throughout dose escalation, we tracked FIREX reporter activities daily. We conducted the experiment with the foresight that reporter fluorescence would enable sorting of resistant cell subpopulations (Fig 3A).

**Figure 3:**
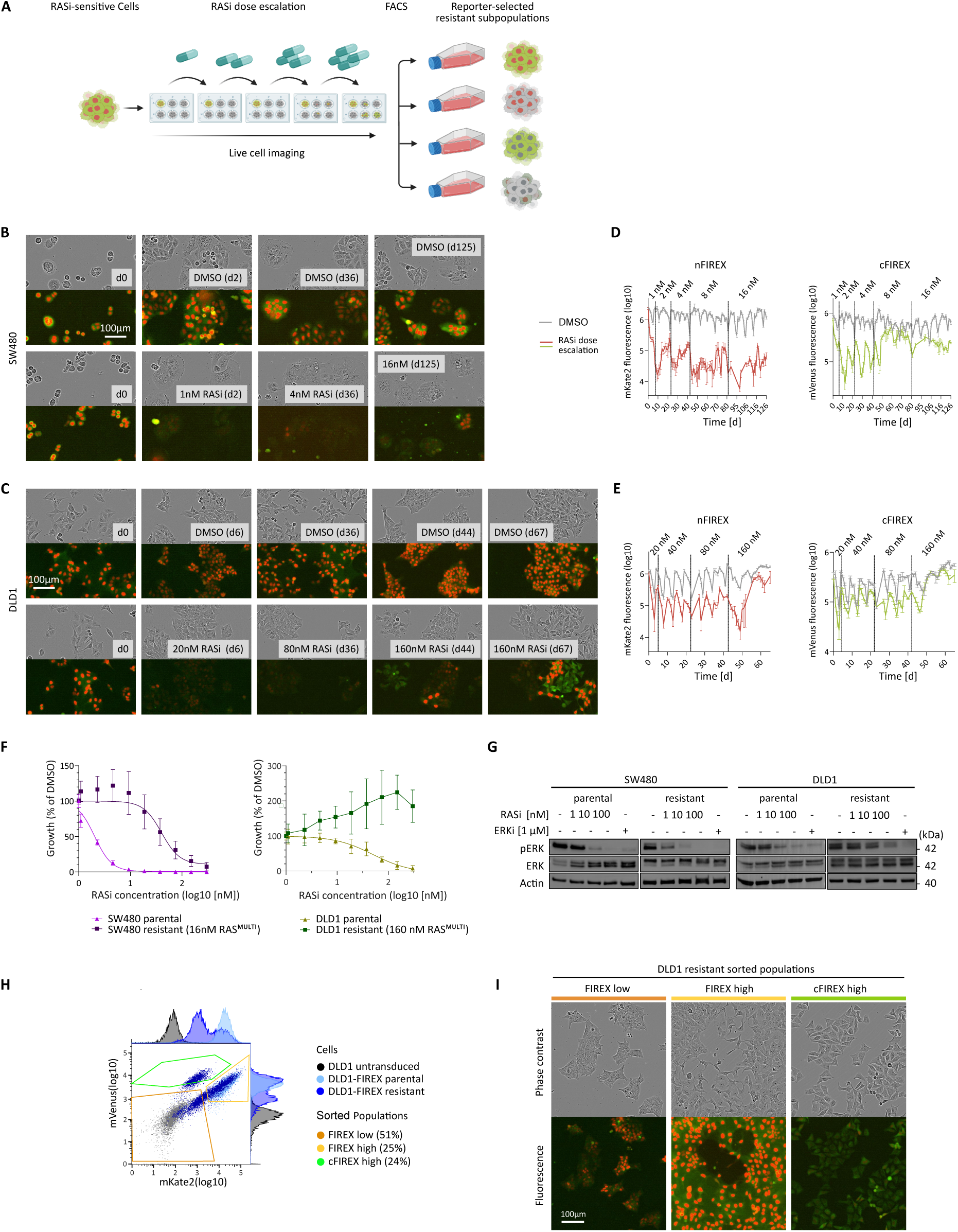
Tracking of ERK activity during development of resistance to RMC-7977 in CRC cell lines. **A** Summary of resistance experiment. In short, stably FIREX-transfected CRC cell lines were subjected to increasing doses of RMC-7977with simultaneous live cell imaging. Resistant subpopulations can be sorted by FIREX pattern to allow individual downstream analysis. **B, C** Tracking of ERK activity during dose escalation in SW480 and DLD1 cells, respectively, at selected time points. Bright-field and fluorescent images are shown. Doses and day after start of treatment are given as insets **D, E** Quantification of FIREX fluorescence over the time course, using RMC-7977 doses as indicated. **F** Cell proliferation in response to RMC-7977 in parental and resistant cell populations, as indicated. **G** ERK phosphorylation in response to RMC-7977 or ERK inhibitor in parental and resistant cell populations, as indicated. **H, I** Selection of RMC-7977-resistant DLD1 subpopulations by fluorescence-activated cell sorting. **H** Final FACS gates for sorting resistant DLD1 cells. For gating strategy, see Suppl. Figure 2). **I** Bright field and fluorescent images of sorted populations.

During dose-escalation, FIREX reporter activities were consistently lower in the cell populations treated with RASi compared to controls (see Fig 3B, C for selected bright field and fluorescent images, and Fig. 3D, E for quantification of reporter activities), although cultures exhibited reporter activity fluctuations during expansion and passaging steps. In SW480 cells, nFIREX remained at low fluorescence levels during selection, whereas cFIREX activity increased by a small factor after approx. 60 days of RASi treatment. In DLD1 cells, distinct subpopulations with varying reporter activities emerged after approximately 40 days of RASi treatment. These subpopulations demonstrated either overall low or high FIREX activities, or predominant cytoplasmic green FIREX activity.

After dose-escalation, we confirmed that the SW480 and DLD1 sublines generated were RASi- resistant (Fig. 3F). The RASi-resistant SW480 cells showed similar levels of ERK phosphorylation in the presence of RASi compared to parental SW480 cells, while the RASi-resistant DLD1 population showed a diminished response of phospho-ERK upon RMC-7977 treatment compared to the parental line (Fig. 3G). To further characterize the resistant DLD1 cells, we used fluorescence-activated cells sorting (FACS) to isolate populations of RASi-resistant DLD1 cells showing generalized FIREX-low or - high, or cFIREX-high reporter patterns, termed DLD1-FIREX low, DLD1-FIREX high or DLD1-cFIREX high, respectively (Fig. 3H, I).

### Validation of reporter-selected differential ERK activity patterns in RASi-resistant DLD1 cells

To validate the distinct ERK activity patterns in the RASi-resistant and FIREX-sorted DLD1 cells, we first screened the ERK-regulated phospho-proteome (*4*) for sites with predicted cell compartment- specificity (Fig. 4A). We shortlisted phospho-EPHA2 (S897), phospho-p90RSK (S380) and phospho- FRA1 (S265) as potential candidates associated with the cell membrane, the cytoplasm and the nucleus, respectively, and verified their subcellular localization by cell fractionation experiments in SW480 and HT29 CRC cells (Fig. 4B). Treatment with RASi and ERKi confirmed the RAS- and ERK- dependence of the phosphorylation sites in CRC cell lines (Fig. 4C).

**Figure 4:**
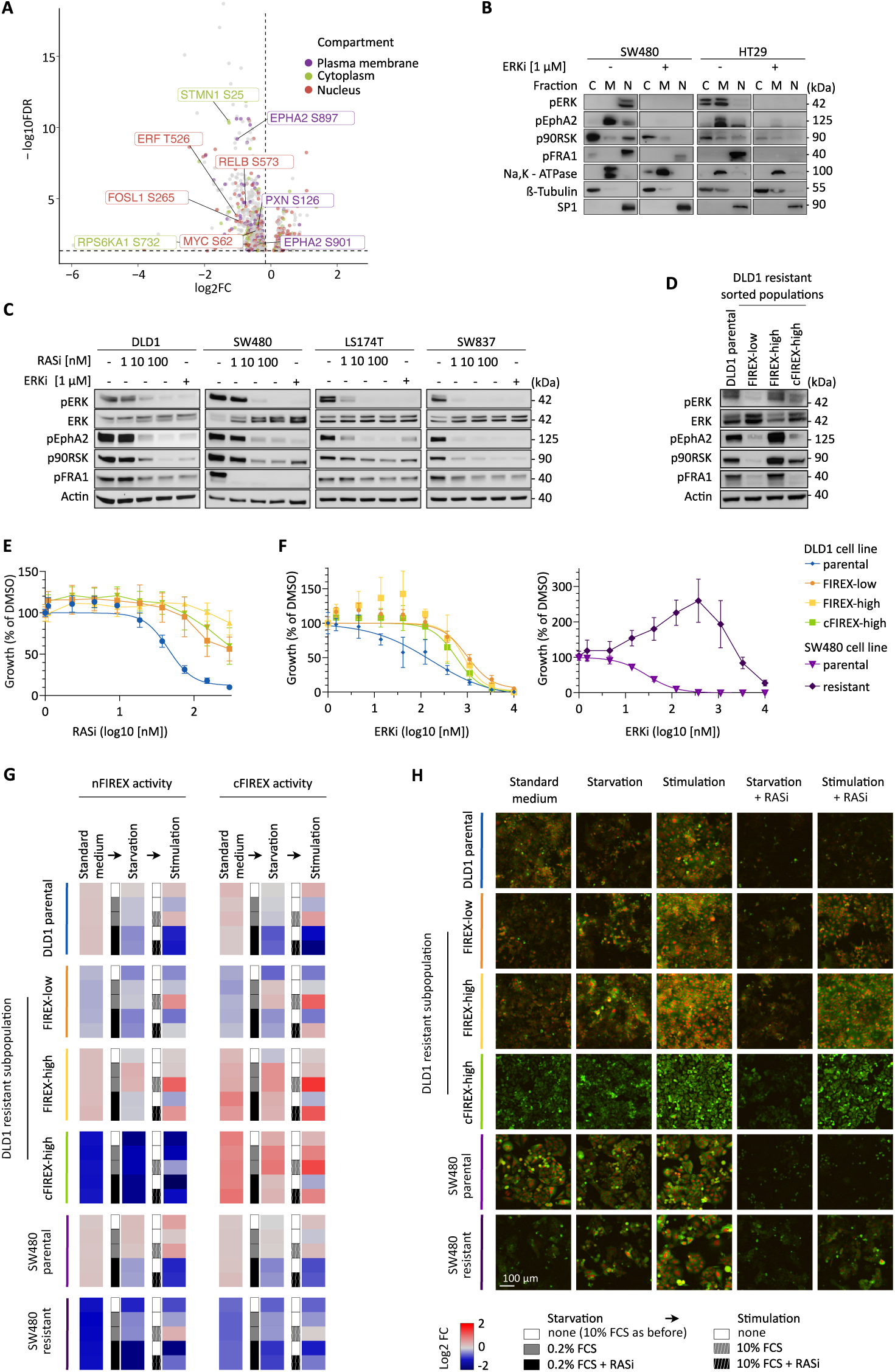
Validation of divergent ERK activity in sorted RASi-resistant DLD1 subpopulations. **A** Identification of ERK targets with potential subcellular-specific localization. Target phospho-sites detected by commercially available antibodies are given. **B** Western blot analyses of subcellular fractions of SW480 and HT29 cells, ± treatment with ERKi for 1 h. pEPHA2, p90RSK and pFRA1 are validated as membranous, cytoplasmic and nuclear ERK targets, respectively. Na,K ATPase, beta-tubulin and SP1 serve as fractionation controls. **C** Response of pEPHA2, p90RSK and pFRA1 to RASi or ERKi in whole cell lysatesfollowing treatment for 1 h. **D** Validation of subcellular divergent ERK activity in RASi-resistant subpopulations by Western blotting of pEPHA2, p90RSK and pFRA1. Resistant cell populations are cultured in the presence of 160nM RASi. **E-F** Cell proliferation in response to RASi or ERKi in parental and resistant cell subpopulations, as indicated, n=3 replicate experiments. **G, H** FIREX reporter activities in parental and RASi-resistant subpopulations, as indicated. **G** Heat map of mean reporter activities before perturbation, 24 h after starvation, and 24 h after re-stimulation, as indicated. Dynamic range normalized to the parental cell lines grown in standard medium. **H** Fluorescent microscopy images of cell culture images at assay endpoint.

Next, we analysed compartment-specific ERK targets in the RASi-resistant DLD1 cell sublines by Western blots (Fig. 4D). In agreement with the ERK reporter patterns, the DLD1-FIREX-low population showed reduced phosphorylation across all ERK-dependent phospho-sites. Conversely, the DLD1- FIREX-high population demonstrated re-activation of all three ERK target sites. Notably, the DLD1- cFIREX high cells sorted for mainly cytoplasmic reporter activity exhibited high activity of the cytoplasmic phospho-p90RSK site, whereas phosphorylation of the nuclear ERK target site on FRA1 remained low.

To ascertain whether the various ERK activity patterns displayed by the RASi-resistant DLD1 cell sublines correspond to distinct reliance on ERK signals, we treated the sublines with RASi or ERKi. Despite the disparities in the phosphorylation of ERK target sites among the sublines, they displayed similar sensitivities to RAS or ERK inhibition (Fig. 4E, F), which were in all cases lower than the sensitivities of the parental cell lines. Similarly, the RASi-resistant SW480 cells also showed reduced sensitivity to ERK inhibition, and these cells proliferated most rapidly when cultured in medium supplemented with 200 nM ERKi (Fig. 4F).

We next quantified FIREX reporter activities in the parental and RASi-resistant cell lines to determine response of ERK to external stimulation in the absence or presence of the RAS inhibitor. For this, we cultured replicates in the presence or absence of serum for 24 h, and subsequently serum-stimulated the starved cells in the absence or presence of RASi for another 24 h (Fig. 4G, H). The perturbation experiment showed that RASi effectively prevented ERK re-activation by serum in the parental DLD1 and SW480 cell lines; however, all resistant cell populations could re-activate the ERK reporter to different degrees following serum starvation and re-stimulation in the presence of RASi. This indicates that all resistant subpopulation have evolved mechanisms that allow for dynamic ERK regulation in the presence of the RAS inhibitor rather than a signalling bypass that acts in parallel to ERK.

### Resistance to broad-spectrum active-state RAS inhibition can be associated with KRAS or RAF1 mutations and specific transcriptome patterns

To identify driver mutations of the RASi-resistant CRC cell subpopulations, we sequenced their exomes at high depth to account for potential cell heterogeneity and called mutations that deviated from the parental cell lines. The resistant DLD1 sublines displayed between 622 and 942 protein- coding mutations at allele frequencies ≥0.01, much more than the 49 mutations found in the RMC- 7977-resistant SW480 cells (Fig. 5A).

**Figure 5:**
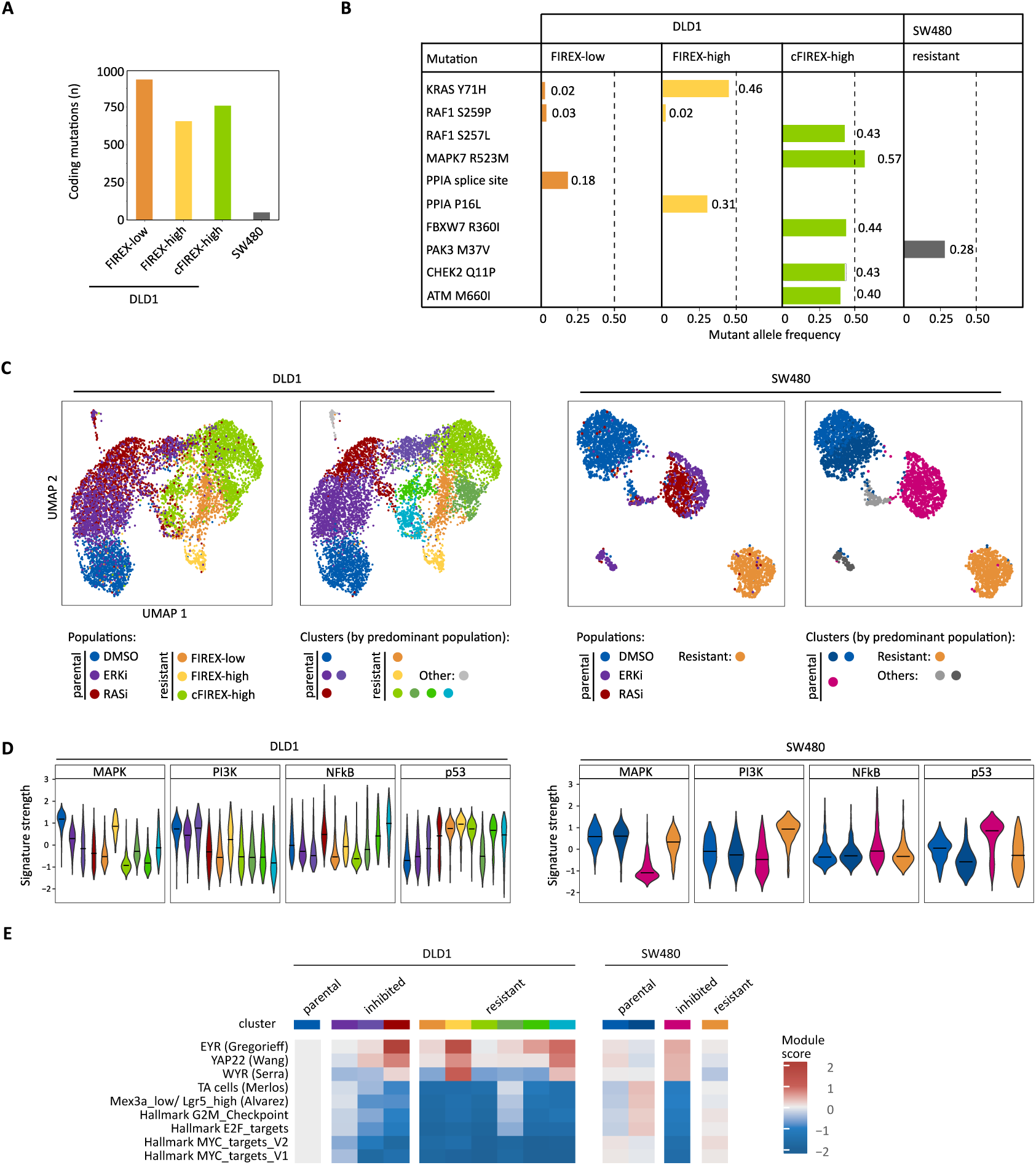
Exome and transcriptome analyses of RMC-7977-resistant cell populations. **A-B** Exome analysis. **A** Load of coding mutations in resistant cell populations, compared to parental cell lines. **B** Putative driver mutations in the resistant cell populations. See supplementary Fig. 5 and Supplementary table 1 for copy number changes in the resistant cell sublines. **C-E** Single cell transcriptome analysis. **C** UMAPs of DLD1 and SW480 cells, as indicated. Analyses encompass parental lines, cultured in the presence or absence of RASi or ERKi, as indicated, and resistant populations cultured in the presence of RASi. UMAPs to the left are color-coded by input samples corresponding to the cell populations, UMAPs to the right indicate clusters, which were color-coded to reflect the predominant cell population. **D** Transcriptional footprints of key signalling pathways, by cluster. See Suppl. Figure 4 for a complete set of signalling pathways. **E** Activities of selected transcriptional signatures relevant in CRC. See Suppl. Figure 5 for activities of the complete list of transcriptional signatures under analysis.

We annotated potential driver mutations using OncoKB (*26*), and in addition explored genes encoding cancer-related signalling proteins (Fig. 5B). The FIREX-sorted and RASi resistant DLD1 subpopulations exhibited minimal overlaps in mutational patterns, thereby validating the sorting strategy. Notably, the DLD1-FIREX high population was defined by a prevalent KRAS Y71H mutation with an allele frequency of 0.46, previously associated with the RASopathy Noonan syndrome and recognized for strengthening RAS-RAF interaction (*27*). Additionally, we identified a low-penetrance RAF1 S259P mutation (*28*), and a splice site mutation within PPIA, encoding the RMC-7977 binding partner Cyclophilin A (*29*). The DLD1-FIREX low population displayed only marginal penetrance of the KRAS and RAF1 mutations (0.02 and 0.03, respectively), but in addition harboured a PPIA P16L mutation. Finally, the DLD1-cFIREX high subpopulation sorted for predominantly cytoplasmic reporter activity was found to have high-penetrance RAF1 S257L (*30*) and MAPK7 R523M mutations. The former mutation is also known to be associated with Noonan syndrome, and the latter is located in the nuclear localisation signal of ERK5 (*31*). Further potential driver mutations exclusive to this cell population include FBXW7 R360I, CHEK2 Q11P, and ATM M660I, impacting an E3 ubiquitin ligase recognized as a colorectal cancer driver and two proteins implicated in cell cycle control, respectively. We could not identify mutations with high probability of acting as genetic drivers in the RASi-resistant SW480 cells; nonetheless we found a PAK3 M37V mutation at an allele frequency of 0.28, affecting a RAF1-regulating kinase (*32*). Analysis of copy-number variations did not identify amplification of KRAS or other key MAPK molecules as potential mechanisms of RAS inhibitor resistance (Supplementary table 1).

To assess transcriptomic footprints associated with RASi resistance, we employed single cell RNA sequencing (scRNA-seq). We compared parental and RASi-resistant sublines, cultured in the absence or presence of RAS inhibitor, respectively. Additional samples of the parental lines were subjected to for RASi or ERKi for 72 h before analysis. This experimental design enables the identification of transcriptomic shifts induced by RAS and ERK inhibition, thereby facilitating assessment of their reactivation upon secondary resistance emergence.

Mostly, the individual cell populations formed distinct clusters, as visualized in a uniform manifold approximation projection (UMAP, Fig. 5C), but DLD1-cFIREX high cells were dispersed across four clusters, indicating transcriptional diversity. In line with the reporter patterns, the DLD1-FIREX high cells exhibited robust reactivation of MAPK target genes, whereas DLD1-FIREX low and DLD1-cFIREX high cells exhibited a lower MAPK transcriptional footprint (Fig. 5D). RASi-resistant SW480 cells exhibited strong expression of MAPK targets, although their FIREX reporter activity was much lower than the parental line. They also had high PI3K-related transcriptional activity.

In addition, the different RASi-resistant CRC populations showed further divergent patterns of gene expression, extending to target genes of NF-kB and p53 (Fig 5D), YAP signalling (*33–35*) or stemness-related gene signatures (*36–38*) (Fig. 5E, Suppl. Fig 4A for a complete list of CRC-related gene expression signatures). It is of note that MYC targets were repressed by RAS or ERK inhibition in both DLD1 and SW480 cells but were re-activated only in RASi -resistant SW480 cells, but not in any of the DLD1 counterparts (Fig. 5E).

### Analysis of RASi-resistant cell exomes, transcriptomes and network states informs combinatorial targeted therapy

Analysis of signalling network states in RAS inhibitor-resistant subpopulations was performed using mass cytometry by time-of-flight (CyTOF). For analysis of steady-state phosphorylation levels, we cultured the parental cells in media without RASi, whereas the resistant DLD1 and SW480 cells were grown in the selection media containing 160 nM and 16nM of RASi, respectively (Fig. 6A, B). Under these conditions, the RASi-resistant populations showed phosphorylation levels of ERK and ERK targets pEPH2 and p90RSK similar to the parental lines, suggesting that they were selected for an optimal level of cytoplasmic MAPK activity(*39*). However, levels of pFRA were strongly reduced in the RASi-resistant DLD1-cFIREX high population with mainly cytoplasmic FIREX activity as well as in the RASi-resistant SW480 cells. The latter observation suggests that the heightened transcription of MAPK targets in SW480 is driven by other factors than FRA1. Concomitant downregulation of vimentin in the RASi-resistant SW480 cells, as also seen in the scRNA-seq data (Suppl. Fig 5B), suggests a potential shift in differentiation state that could command heightened expression of MAPK targets.

**Figure 6:**
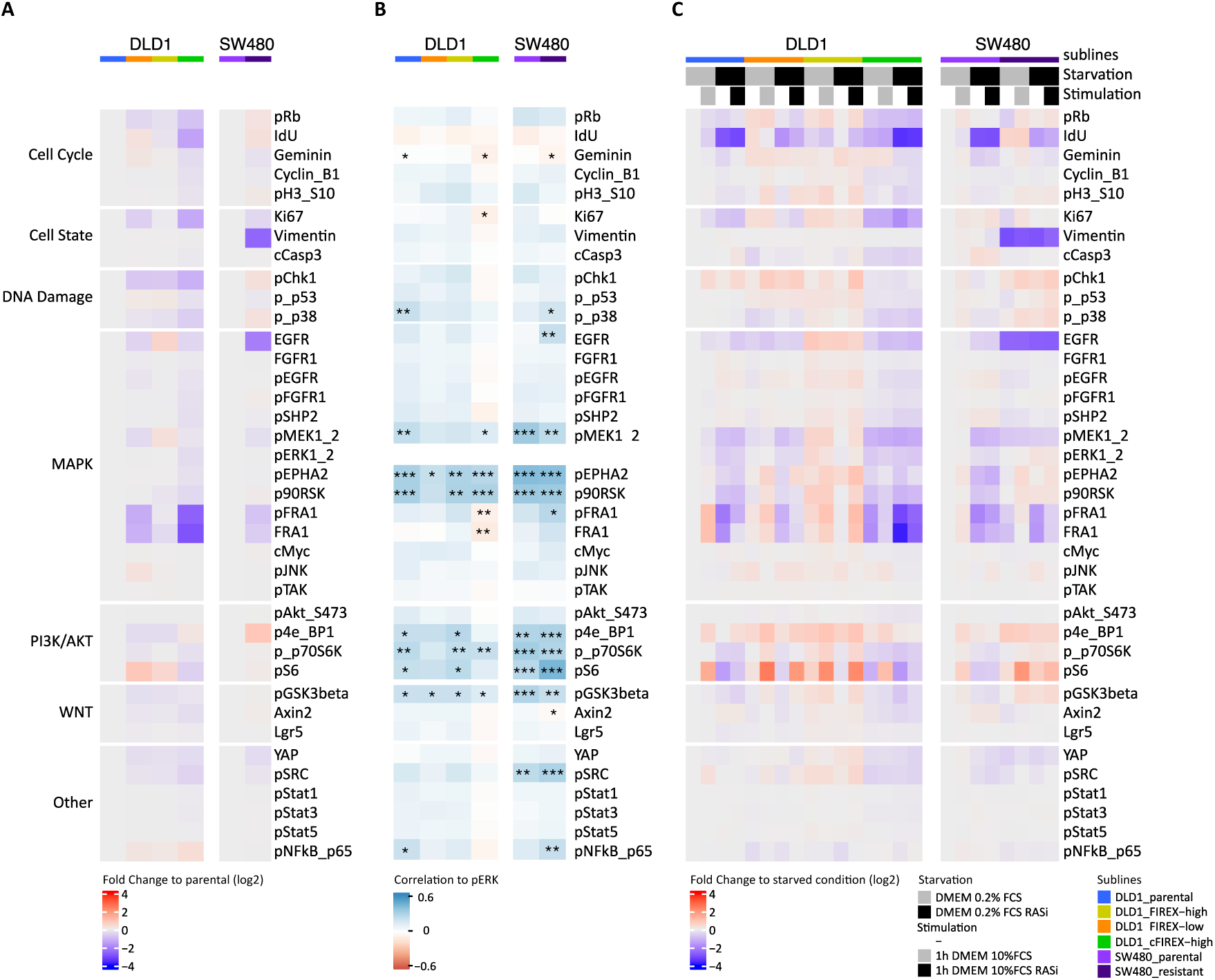
Mass Cytometry reveals adaptations in cell signalling networks upon development of RMS-7977 resistance. **A** Average activities of cell cycle, cell states, DNA damage markers and signalling network nodes in parental versus resistant CRC cell lines, as indicated. **B** Correlations of markers with ERK phosphorylation, calculated from data in A. **C** Network perturbation analysis of parental versus resistant CRC cell lines, before and after serum starvation, serum re-stimulation and RMC-7977 treatment, as indicated. Median of n=3 biological replicates. See Suppl. Fig 6. for data of each replicate and complete correlation maps.

On a cell-to-cell basis, we found strong positive correlations between phosphorylation of ERK and further signalling nodes, thereby defining an ERK-centred signaling network (Fig. 6B). This included the cytoplasmic ERK targets pEPHA2, and p90RSK and extended to PI3K-AKT signal transducers such as 4e-BP1, p70S6K and S6, suggesting that in general the RAS downstream MAPK and PI3K-AKT pathways are activated in parallel across all parental and resistant CRC cell populations under analysis. However, phosphorylation of FRA1 was decoupled from the ERK network in the RASi- resistant DLD1-cFIREX high line with cytoplasmic FIREX activity, corroborating the Western blot and single cell transcriptome analyses.

Network perturbation by serum starvation and re-stimulation in the presence and absence of RASi furthermore showed highest MAPK network dynamics in the RASi-resistant DLD1-FIREX high cells compared to the other parental and resistant sublines (Fig. 6C), whereas the AKT-PI3K axis showed increased dynamics in multiple RASi-resistant populations when compared to the parental lines. In agreement with the ERK reporter assays (Fig. 4G, H, above), RASi prevented serum-induced MAKP and PI3K-AKT signalling in the parental lines, but not in the resistant derivatives.

Our exome, transcriptome and phospho-protein analyses revealed key differences between the RASi-resistant CRC cell populations (Fig. 7A). We finally asked whether these can be exploited to restore RASi-sensitivity in the resistant cell populations. As potential vulnerabilities we chose RAF, as it is mutated in the cFIREX high DLD1 subpopulation (Fig. 5B, above), EGFR as a key player of MAPK feedback signalling downregulated in multiple RASi-resistant subpopulations (Fig. 6A, above), and AKT, as multiple signal transducers of the PI3K-AKT axis are differentially regulated between the resistant DLD1 subpopulations (Fig. 6A, C, above). Subsequent investigations involved assessing the sensitivity of parental cell lines and RASi-resistant populations to Naporafenib (LXH254), Sapitinib (AZD8931), and MK-2206, serving as inhibitors of RAF, EGFR, and AKT, respectively, in conjunction with RASi (Fig. 7B-D). While most inhibitor combinations showed no combinatorial effects, the RAF inhibitor Naporafenib showed strong synergy with RASi specifically in the DLD1 cFIREX-high cell population harboring the RAF1 S257L mutation (Fig. 7B, C). This discovery implies the potential of RAS-RAF inhibitor combinations to mitigate clonal expansion within RAF1-mutant and RASi-resistant cell populations.

**Figure 7:**
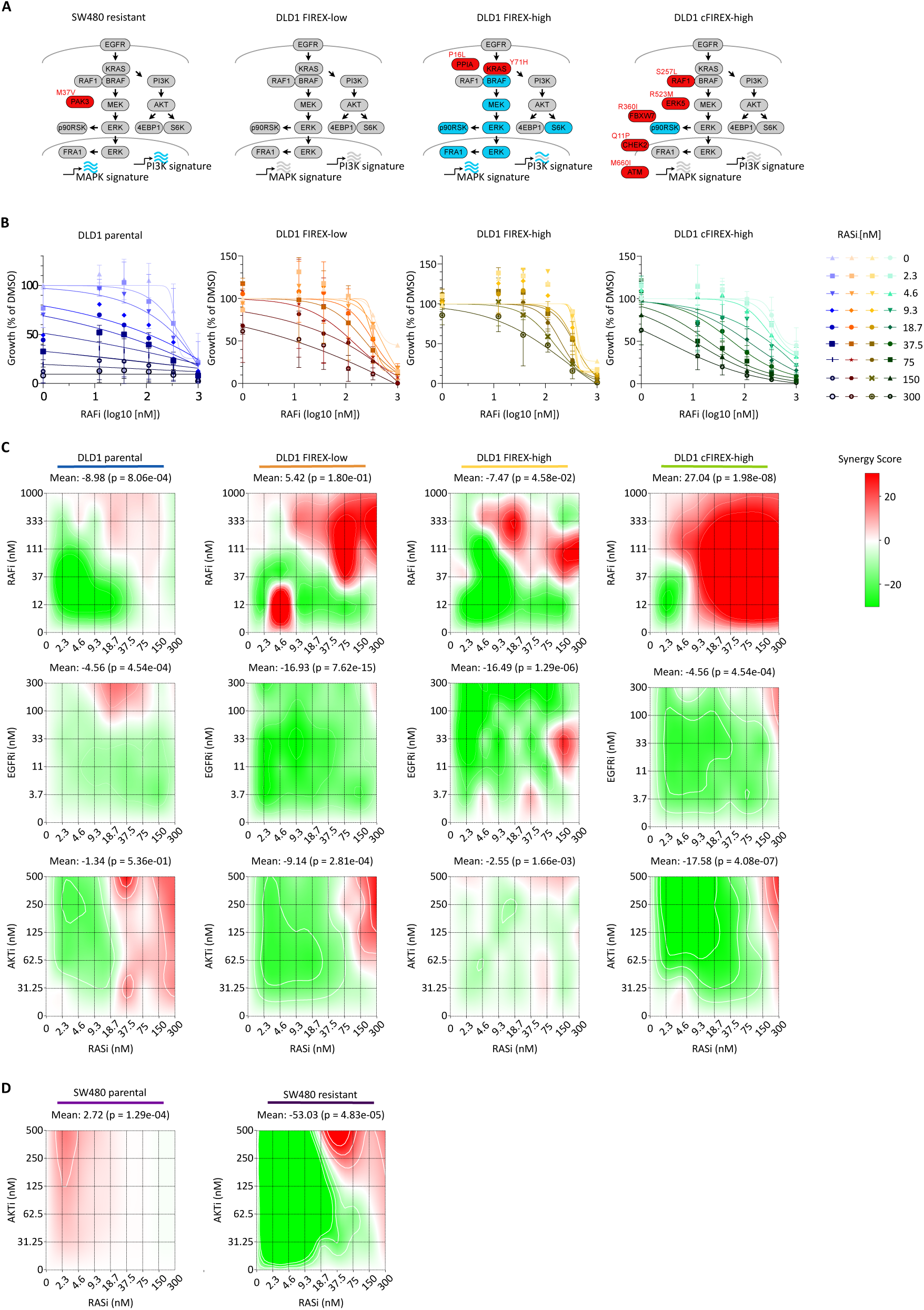
RASi-resistant cell exomes, transcriptome and network states inform combinatorial targeted therapy. **A** Cell signalling network states in the RASi-resistant cell populations, as inferred by exome, mass cytometry and transcriptome analyses. Grey: low activity, blue: high and dynamic activity; red: mutated **B-D** Combinatorial drug treatments of subpopulations, as indicated, mean of n=3 biological replicates. **B** Response of parental and RMC-7977-resistant DLD1 cell populations to combinatorial RAS and RAF inhibition. **C, D** Synergy maps showing response of parental and resistant cell populations to RMC-7977 and RAF inhibitor Naporafenib (LXH254), EGFR inhibitor Sapitinib (AZD8931), or AKT inhibitor MK-2206, as indicated. **C** DLD1 cell populations, **D** SW480 cell populations.

## Discussion

Multiple studies have probed the mechanisms underpinning resistance to mutation-specific inactive-state inhibitors of RAS (*14, 17, 18, 40*), but studies on resistance to broad-spectrum active-state RAS inhibition are only now emerging. Focussing on CRC, we found that RAS-mutant cell lines are universally sensitive to the active-state RAS inhibitor RMC-7977. Using a RASi dose escalation experimental design (Fig. 3), we observed the emergence of multiple populations of inhibitor-resistant cells with distinct patterns of ERK reporter reactivation that allowed for cell sorting. We found that all four RASi-resistant cell populations of two CRC cell lines had developed differential mechanisms enabling dynamic ERK regulation when cultured in the presence of the RAS inhibitor (Fig. 4G, H and Fig. 6C), as opposed to relying on bypass mechanisms that typically result in resistance without ERK reactivation. Exome sequencing identified a KRAS Y71H mutation in cells demonstrating strong MAPK pathway reactivation. Furthermore, we identified a RAF1 S257L mutation in a RASi-resistant cell population with lower level and cytoplasm-centred of MAPK pathway reactivation.

These KRAS and RAF1 mutations are rare in cancer but have previously been identified in the heritable RASopathy Noonan syndrome. We found that the RASi-resistant cell population with a prevalent RAF1 S257L mutation is sensitive to concurrent RAS and RAF inhibition. This finding can serve as a starting point for exploring combinatorial treatments aimed at mitigating resistance to active-state RAS inhibition in clinical settings.

Noonan syndrome, an autosomal dominant disorder resulting in congenital heart defects and multiple characteristic developmental features, arises from somatic mutations in the RAS-MAPK pathway (*41*). The KRAS Y71H, RAF1 S257L and RAF1 S259P mutations identified here as putative drivers of active-state RAS-inhibitor resistance have all been described in Noonan syndrome patients, implying that they are generally non-oncogenic on their own across somatic tissue types. Molecular characterisation of KRAS Y71H has revealed increased complex formation with RAF1 compared to wildtype KRAS (*27*). Mutations of RAF1 at codons 257 or 259 are known to prevent 14-3-3 protein binding and stabilization of an autoinhibited RAF1 state, thus likewise favouring RAS-RAF interaction (*42, 43*). Therefore, it is tempting to speculate that the mutations we identified compete with the assembly of the trimeric RAS-Cyclophilin A-RMC-7977 complex by favouring formation of the alternative RAS-RAF complex. It is noteworthy that this mechanism is distinct from the hotspot point mutations driving resistance to mutation-specific KRAS G12C/D inhibitors that have oncogenic activity on their own.

In our exome sequencing, we also find two mutations in PPIA, encoding Cyclophilin A. Although the functional significance of the mutations is unclear, their presence suggests that the trimeric inhibitor complex could also serve as a structural target for the emergence of RAS inhibitor resistance.

The effects of active ERK are highly complex, with a plethora of cytoplasmic and nuclear targets (*4, 44–47*). Various ERK reporters excel at capturing individual aspects of ERK activity (*46*), however no design was able to integrate nuclear and cytoplasmic ERK activities independently. Here, we provide a second-generation ERK reporter system tailored for temporal integration of ERK activity (*23*), while providing distinct fluorescent readouts for the nucleus and the cytoplasm. The reporter system allowed us to sort individual subpopulations of RASi-resistant DLD1 cells with unique mutational profiles and phospho-protein network dynamics. In contrast to the parental cells, all RASi-resistant populations could activate the ERK reporter after serum starvation and restimulation in the presence of RMC-7977, albeit to different degrees as visualized by the reporter (Fig 4G, H) and also confirmed by mass cytometry analysis (Fig. 6C). This finding indicates that resistance generally arises from alterations within the MAPK pathway, and that a KRAS Y71H mutations allows for higher network dynamics towards external stimuli than alternative RASi-resistance driver mutations such as in RAF1.

Reactivation of ERK was confined to the cytoplasm in the sorted RASi-resistant DLD1-cFIREX high subpopulations, as indicated by reporter fluorescence (Fig. 3I) and Western blot analysis (Fig. 4D) alike. Mass cytometry revealed high correlation between phosphorylation levels of ERK and its cytoplasmic target p90RSK but no such link to the nuclear ERK target FRA1 in these cells (Fig. 6B). In addition, the scRNA-seq data revealed low transcriptional activity of MAPK and MYC target gene signatures. Thus, multiple assays indicate decoupling of cytoplasmic from nuclear ERK activity in the DLD1-cFIREX high cells. As FRA1 and MYC were recently highlighted as crucial nuclear executers of oncogenic RAS-MAPK signals (*45*), it remains to be determined how reactivation of cytoplasmic ERK activity alone can drive resistance to RASi. Our study identified multiple potential drivers with high penetrance, including RAF1 S257L, ERK5 R523M, FBXW7 R360I, CHEK2 Q11P, and ATM M660I which may take part in establishing RASi resistance in the absence of MAPK target gene activation.

However, our finding of mainly cytoplasmic ERK re-activation during emergence of secondary resistance to RAS-MAPK inhibition is not unprecedented, as resistance of melanoma to BRAF inhibition was also found to be correlated to cytoplasmic, but not nuclear, ERK activity (*48*). Generally, the equilibrium between cytoplasmic and nuclear ERK activity can be influenced by dual-specificity phosphatases (*49*), attachment of ERK to cytoplasmic scaffolds (*50, 51*), or different modes of transportation (*52–54*), and future studies are required to explore these mechanisms in the emergence of secondary resistance during therapeutic RAS-MAPK inhibition.

In RASi-resistant SW480 cells, we observe diminished ERK reporter activity and low presence of FRA1 in mass cytometry, but reactivation of MAPK, PI3K and MYC gene expression signatures (Fig. 5D, E). Together, the disconnect between the signalling pathway activities and transcriptional footprints may imply epigenetic re-programming. As we see a strong and consistent downregulation of vimentin in the resistant SW480 cells compared to the parental line in our single cell transcriptome and mass cytometry data, we hypothesize that these cells may have undergone a histological transformation that is known to underly many clinical resistances to anti-MAPK therapy in the clinic (*55*) and in experimental models (*56*).

In summary, our study serves as a proof-of-concept for reporter-assisted cell sorting to disentangle resistance mechanisms by observing pathway reactivation in real time. Integration of the ERK reporter system with a larger panel of cell lines and an expanded set of inhibitor combinations could yield a more comprehensive map of resistance drivers and combinatorial treatment options for broad-spectrum active-state RAS inhibition. The approach offers the capability to monitor clinically relevant resistances within clonal populations as they evolve, thereby enhancing our understanding of resistance mechanisms and facilitating the exploration of therapeutic interventions.

## Materials and Methods

### Cell culture

Cells were cultured in Dubeccós Modified Eaglés medium (DMEM, Lonza, BE12-707F) supplemented with 10% fetal calf serum (Pan-Biotech, P30-1506), except when noted otherwise.

For lentiviral production, HEK293T cells were transfected with viral vectors using polyethylenimine (PEI, MedChemExpress, HY-K2014) reagent, using the packaging plasmids PsPAX2 and PRSV-Rev, along with 2 µg of the envelope plasmid pMD2.G. The next day, the medium was changed to allow for cell recovery. Over the subsequent two days, supernatant was collected twice and stored at 4°C. The collected supernatant was then filtered through a 0.45 µm filter and concentrated using LentiX concentrator (TaKaRa, 631232). For long-term storage, the concentrated virus was aliquoted and stored at -80°C.

The following inhibitors were used in the study:

| Inhibitor | Primary Target | Manufacturer | Product No. |
| --- | --- | --- | --- |
| Adagrasib (MRTX849) | KRAS G12C | Selleck Chemicals | S8884 |
| Encorafenib | BRAF V600E | Selleck Chemicals | S7108 |
| Fadraciclib (CYC065) | CDK2/9 | Selleck Chemicals | S7371 |
| JNK-IN-8 | JNK | MedChemExpress | HY-13319 |
| MK-2206 | AKT | Selleck Chemicals | S1078 |
| MRTX1133 | KRAS G12D | Selleck Chemicals | E1051 |
| Naporafenib (LHX254) | RAF | MedChemExpress | HY-112089 |
| Palbociclib | CDK4/6 | Selleck Chemicals | S4482 |
| RMC-7977 | RAS | Revolution Medicines |  |
| Sapitinib (AZD8931) | EGFR | Selleck Chemicals | S2192 |
| SB203580 | p38 | Selleck Chemicals | S1076 |
| SCH772984 | ERK1/2 | Selleck Chemicals | S7101 |
| Trametinib | MEK1/2 | Selleck Chemicals | S2673 |

For generation of RMC-7977-resistant CRC cells, cells were seeded at a density of 1 x 10^5^ cells per well in 1 ml of cell culture medium containing DMSO or RMC-7977 (Revolution Medicines).

Logarithmically growing cells were initially treated with GI-30 dosages of the inhibitor in the respective cell line and concentrations were doubled, once cells resumed proliferation similar to the parental line.

### Reporter generation

cFIREX and nFIREX were created by generating mKate2, cMyc NLS and IkBa NES sequences from via gene synthesis (Eurofins), and employing mVenus and Fra1 163-271 sequences from the original FIRE reporter (*23*). Synthesis products were amplified by PCR and assembled by T4 ligase (Promega, C671B), using NheI and PacI sites. Final plasmids were sequenced to confirm integrity. Plasmids and sequences are available by request.

### Reporter cell line generation

For generation of stable FIREX CRC cell lines, cells were seeded at 50% confluency. Concentrated nFIREX and cFIREX virus was diluted at a ratio of 1:5 with media, and 8 µg/ml of polybrene was added to enhance transduction efficiency. After transduction of both nFIREX and cFIREX, cells were subjected to fluorescence-activated cell sorting and gated for intermediate to high red and green fluorescence. Cells were individually isolated into wells of 96-well plates and expanded. Monoclonal sublines underwent dynamic response testing using live-cell imaging in Incucyte using either DMSO or 100 nM MEKi (Trametinib, Selleck Chemicals, S2673), and sublines exhibiting the strongest dynamic response were chosen for further experimentation. Sublines were split once they reached 70% confluency, and an equal cell count was seeded. Upon reaching proliferation rates comparable to the control cell line, the drug concentration was increased using a stepwise dose escalation strategy to generate stable resistant sublines. Furthermore, FIREX-transduced cell lines (DLD1 and SW480) were subjected to daily observation through live-cell imaging in an Incucyte cell culture microscope (Sartorius).

To evaluate the dynamic response of the reporters, HEK293 RAF:ER cells transduced with FIRE, cFIRE, cFIREX, and nFIREX were either starved in medium containing 0.2 % FCS for 24 hours or stimulated with 5 µM 4-Hydroxy-Tamoxifen (4OHT, Sigma Aldrich, H7904).

### Flow cytometry and fluorescence-activated cell sorting (FACS)

For flow cytometry, cells were harvested using TrypLE (Gibco 12604-013), resuspended in PBS, and filtered through 30 µm Celltrics filters (Sysmex) and recorded using a CytoFLEX S (Beckmann Coulter) flow cytometer. For mVenus detection, a 488 nm laser with a 525/40 bandpass filter was employed, while for mKate2 detection, a 638 nm laser with a 610/20 bandpass filter was used. In addition, the untransduced cells were examined to establish gating parameters and determine the proportion of cells lacking transduction. FCS files were analyzed using FlowJo software (Becton Dickinson).

CRC cells transduced with FIREX were sorted using a FACSMelody Cell Sorter (Becton Dickinson). Prior to sorting, cells were harvested using TrypLE, resuspended in medium, and filtered through 30 µm CellTrics filters. Untransfected cells, and cells containing single cFIREX or nFIREX reporter plasmids were collected to calibrate the gates and apply automated spillover matrix calculation. For detection of mVenus and mKate2 expression levels, FITC 488 nm with 527/32 bandpass filter and PE 561 nm with 613/18 bandpass filter were employed, respectively. Spillover matrix calculation was applied to samples during the run. Subsequently, gates of interest were defined and cell lines were sorted either into tubes or 96-well plates, generating polyclonal or monoclonal colonies, respectively.

### Proliferation and apoptosis assays

To determine cell proliferation, cell numbers were quantified by propidium iodide staining of nuclei. For this, cells were seeded in triplicates into 96-well plates at 1500 cells/well. After 24 hours, drugs were added using a D300e Digital Dispenser (Tecan). Before counting, plates were fixed with cold 70% ethanol and stored at -20°C. For staining, plates were washed with PBS and 10 μg/ml propidium iodide (ThermoFisher, BMS500PI) was added. Staining proceeded for 30 minutes at room temperature in the dark. Subsequently, the plates were washed once with PBS before imaging. PI fluorescence was was measured using a 561 nm laser and detected through a 610 nm bandpass filter using a CQ1 confocal fluorescence microscope (Yokogawa). Nuclei were counted using CellPathfinder (Yokogawa). Data were analysed using Prism software (GraphPad). Synergy scores were computed using synergyfinder(*57*), applying a drug additivity model developed by Loewe.

For apoptosis assays, 3-5 x 10^5^ cells were seeded per well in 6-well plates. After 24 hours, the medium was refreshed, and drugs were administered as specified. Additionally, as a positive control for apoptosis, cells were treated with 1 µM Staurosporine (STS, Merck, 569397). Following 48 hours of treatment, both floating and adherent cells were harvested, washed twice with PBS, and harvested by centrifugation. Next, cells were fixed with 2% formaldehyde for 10 minutes at RT. Samples were washed with PBS and permeabilized with pre-chilled 90% methanol on ice for 30 minutes before being stored at -20°C. For analysis, the samples were washed twice with incubation buffer (0.5% BSA in PBS) and stained with an anti-cleaved caspase-3 Alexa Fluor® 488 antibody (Cell Signaling, 9603, 1:50) for 1 hour at room temperature. Following staining, the cells were washed with incubation buffer, resuspended in PBS, and analyzed using the CytoFLEX S flow cytometer. Data obtained were analyzed using FlowJo software (FlowJo, LLC).

### Live cell imaging

For analysis of individual cells, the CQ1 microscope (Yokogawa) and CellPathfinder software (Yokogawa) was used. For population analyses, the Incucyte cell culture microscope (Sartorius) was used. Green and red fluorescence channels were acquired for 500 and 400 ms, respectively, followed by spectral unmixing of 10% of the red channel towards the green. Subsequently, the images were analyzed using the Incucyte software in Basic Analyzer mode, wherein cell, cytoplasm, and nucleus were defined. Finally, the data on the integrated fluorescence of green or red per image, normalized to the cell area in the image, was exported, and graphs were generated using Prism (GraphPad).

### Western blot analysis

For western blots, cells were harvested, collected by centrifugation and lysed in MPER/RIPA buffer (ThermoFisher, 89900) combined with PhosSTOP (Sigma, 4906837001) and Complete protease inhibitor (Sigma, 4693159001) cocktail. Lysates underwent 5 minutes of sonication with on/off intervals of 30 seconds using a Bioruptor (Diagenode), followed by centrifugation at 20,000 g for 10 minutes to retrieve the protein supernatant. Cell fractionation was performed using the subcellular protein fractionation kit for cultured cells (ThermoFisher 78840), following the manufacturer’s protocol. Protein concentration was assessed using a BCA protein quantification kit, according to the manufacturer’s guidelines. 25 µg of protein lysates were combined with 6xSDS sample buffer and boiled for 5 minutes at 95°C with agitation. After cooling, the samples were loaded onto 10% TGX Stain-Free polyacrylamide gels for SDS-PAGE. Proteins were transferred to a PVDF membrane using tank blotting at 80 V for 90 minutes. Subsequently, the membranes were blocked with 5% milk TBS-T blocking buffer for 1 hour at room temperature or overnight at 4 °C, followed by washing with TBS-T. Membranes were incubated with the respective primary antibody diluted in blocking buffer (see Table 7) for 2 hours at room temperature or overnight at 4°C. After washing three times with TBS-T for 5 minutes each, membranes were incubated with a secondary antibody in blocking buffer for 1 hour at room temperature. Protein detection was performed using Clarity Western ECL (Biorad) or SuperSignal West Femto Substrates (Thermofisher) and a ChemiDoc MP Imaging System (Qiagen).

The following primary antibodies were employed:

| Target | Vendor | Product No. | Usage |
| --- | --- | --- | --- |
| Actin (Rhodamine) | BioRad | 12004164 | 1:3000 |
| Cleaved Caspase-3 (Asp175) (D3E9)<br>Rabbit mAb (Alexa Fluor® 488 Conjugate) | Cell Signaling | 9603 | 1:50<br>Flow Cytometry |
| ERF (Thr526) | Cohesion Biosciences | CPA4404 | 1:500 |
| ERK1/2 p42/44 | Cell Signaling | 4696 | 1:1000 |
| FLNB (Ser2197) | Cell Signaling | 12005 | 1:1000 |
| FRA1 | Cell Signaling | 5281 | 1:500 |
| LCP1 (Ser5) | ThermoFisher Scientific ` | AB_2817224 | 1:500 |
| MYC (Ser62) | ThermoFisher Scientific ` | PA5-118565 | 1:500 |
| Na,K-ATPase | Cell Signaling | 3010 | 1:1000 |
| p90RSK (Ser380) | Cell Signaling | 12032 | 1:500 |
| p-c-Jun (Ser73) | Cell Signaling | 3270 | 1:1000 |
| pEphA2 (Ser897) | Cell Signaling | 6347 | 1:500 |
| pERK1/2 p42/44 (Thr202/Tyr204) | Cell Signaling | 9101 | 1:1000 |
| pFRA1 (Ser265) | Cell Signaling | 5841 | 1:500 |
| pMAPKAPK-2 (Thr334) | Cell Signaling | 3007 | 1:1000 |
| PPP1R12A (Ser507) | Cell Signaling | 3040 | 1:500 |
| pRb (Ser807/811) | Cell Signaling | 9308 | 1:1000 |
| pRpb1 CTD (Ser2) | Cell Signaling | 13499 | 1:1000 |
| RAF1 (Ser642) | Cell signalling | 12005 | 1:1000 |
| RELB (Ser573) | MyBioSource | MBS9402624 | 1:500 |
| REPS1 (Ser709) | MyBioSource | MBS9207978 | 1:1000 |
| SP1 | Abcam | ab13370 | 1:2500 |
| STMN1 (Ser25) | Cohesion Biosciences | CPA4529 | 1:500 |
| STMN1(Ser16) | Abcam | ab47328 | 1:500 |
| β-Tubulin | NEB | 2146 | 1:1000 |

### Mass cytometry

A published CyTOF protocol (*58*) was used with modifications. In short cells were incubated for 30 minutes with 5 µM Cell-ID IdU (Fluidigm, 201127) before harvesting, washed with PBS, and digested to a single cell solution using TrypLE supplemented with 100 U/ml Universal Nuclease (ThermoFisher, 88701) at 37°C. Subsequently, cells were counted, and a maximum of 5 x 10^5^ cells were stained with 2.5 μM Cisplatin (Fluidigm, 201064) in MaxPar PBS (Fluidigm, 201058) for 5 minutes at 37°C. Subsequently, cells were resuspended in their respective growth medium and allowed to rest for 5 minutes at 37°C. Cells were resuspended in a 10% BSA-PBS solution, mixed 1:1.4 with Proteomics Stabilizer (SmartTube, PROT-1), and incubated for 10 minutes at room temperature and frozen at - 80°C for storage.

For barcoding and antibody staining, cells were thawed in a 37 °C water bath, MaxPar Cell Staining Buffer (Fluidigm 201068) was added, and supernatant was removed. Cell pellets underwent washing in Cell Staining Buffer before being resuspended in Barcode Perm Buffer (Fluidigm, 201057).

Barcodes (Fluidigm, S00114) in Barcode Perm Buffer were mixed with the sample and incubated for 30 minutes at room temperature, before washing with Cell Staining Buffer and pooling. Pooled cells were filtered, permeabilized with cooled methanol, washed and incubated with the antibody cocktail for 30 minutes, followed by incubation with the 62.5 nM Cell-ID Intercalator-Ir. After overnight fixation in methanol-free formaldehyde (2% in PBS, Pierce, 28906) at 4°C, final washes were performed before storing the cells on ice until measurement. CyTOF acquisition was conducted on a HELIOS machine (Fluidigm). FCS files were analysed using R. Unwanted covariance was then mitigated using the R package RUCova (*59*).

The following CyTOF antibodies were used:

| Target | Concentration | Vendor | Clone | Product No. |
| --- | --- | --- | --- | --- |
| Axin 2 | 2 µg/ml | Abcam | polyclonal | ab32197 |
| cCasp3 | 1 µl from stock per sample | Fluidigm | D3E9 | 3142004A |
| Cyclin B1 | 2 µg/ml | BD | GNS-11 | 554178 |
| EGFR | 2 µg/ml | Novus | 31G7 | NBP2-47740 |
| FGFR1 | 5 µg/ml | CST | D8E4 | 9740BF |
| FRA1 | 2 µg/ml | Abcam | EPR23767-177 | ab278103 |
| Geminin | 4 µg/ml | Proteintech |  | 10802-1-AP |
| Ki-67 | 1 µl from stock per | Fluidigm | B56 | 3162012B |
| LGR5 | 1 µl from stock per | Fluidigm | 4D11F8 | 3161025B |
| MYC | 1 µl from stock per | Fluidigm | 9.00E+10 | 3176012B |
| p-FGFR1 | 6 µg/ml | CST | D4X3D | 52928BF |
| p-GSK3β [S9] | 4 µg/ml | CST | D85E12 | 79774SF |
| p-JNK | 4 µg/ml | CST | 81E11 | ab236152 |
| p-p38 [T180/Y182] | 1 µl from stock per | Fluidigm | D3F9 | 3156002A |
| p-p53 [S15] | 2 µg/ml | CST | 16G8 | 56979 |
| p-p70S6K [T389] | 4 µg/ml | CST | 1A5 | 9206 |
| p-p90RSK [S380] | 4 µg/ml | CST | D5D8 | 12032 |
| p-SRC [T418] | 2 µg/ml | eBioscience | 14-9034-82 | SC1T2M3 |
| p-STAT1 | 2 µg/ml | BioLegend | A17012A | 666402 |
| p-STAT5 | 4 µg/ml | Fluidigm | D47E7 | 4322 |
| p4e-BP1 [T37/46] | 2 µg/ml | CST | 236B4 | 2855BF |
| pAkt [S473] | 1 µl from stock per sample | Fluidigm | D9E | 3152005A |
| pan Akt | 4 µg/ml | CST | 40D4 | 28544SF |
| pChk1 [S345] | 4 µg/ml | CST | 133D3 | 76784SF |
| pEGFR [Y1068] | 1 µl from stock per sample | Fluidigm | D7A5 | 3146007A |
| pEphA2 | 5 µg/ml | CST | D9A1 | 6347BF |
| pERK1/2 [T202/Y204] | 1 µl from stock per | Fluidigm | D13.14.4E | 3171010A |
| pFRA1 | 20 µg/ml | CST | D22B1 | 5841BF |
| pH3 [S10] | 1 µg/ml | CST | D2C8 | 3377 |
| pMEK1/2 [S217/221] | 2 µg/ml | CST | 41G9 | 9154BF |
| pNF-κB p65 [S536] | 2 µg/ml | CST | 93H1 | 3033BF |
| pRb [S807/S811] | 1 µg/ml | BD | J112-906 | 558389 |
| pS6 [S235/236] | 1 µl from stock per | Fluidigm | N7-548 | 3175009A |
| pSHP2 | 1 µl from stock per sample | Fluidigm | D66F10 | 3141002A |
| pSTAT3 [Y705] | 1 µl from stock per | Fluidigm | 4/P-STAT3 | 3158005A |
| pTAK1 [S412] | 4 µg/ml | Bios | polyclonal | BS-3435R |
| total Erk1/2 | 4 µg/ml | CST | L34F12 | 78241BF |
| Vimentin | 4 µg/ml | CST | D21H3 | 5741BF |
| YAP | 2 µg/ml | CST | D8H1X | 14074BF |

### Identification of compartment-specific ERK targets

To define compartment-specific ERK targets, we re-analyzed the published data (PRIDE accession ID PXD048529) (*4*) of six pancreatic cell lines treated with ERK inhibitor SCH772984 or solvent control for 1 h or 24 h. Data was filtered for sufficient localization probability (>0.75), log2-transformed and normalized by sample-wise median-centering. We then calculated log2 ratios to respective mean DMSO control. To derive a list of ERK significant consensus targets we applied the R package *limma* (*60*) on the log2 ratios using factors for treatment and time point and blocking information of cell lines. Next, we selected candidate sites by the following criteria: (1) significant downregulation upon 1 h ERK inhibition (FDR <=0.05), (2) persistent downregulation after 24 h ERK inhibition and (3) exclusive compartmentation as cytoplasmic, nuclear or plasma membrane (using GO cellular localization criterion from R package *GO.db*). The remaining 200 phospho sites were then investigated for available antibodies.

### Single cell RNA sequencing

For single cell RNA sequencing, cells were harvested using TrypLE, resuspended in PBS, filtered through 30 µm Celltrix filters, resuspended in cell freezing medium and stored at -80°C until further processing. Cryopreserved cells were quickly thawed in a 37°C waterbath, washed in cell culture medium and resuspended in 1xPBS supplemented with 10% fetal calf serum. Cells were counted using an automated cell counter TC20 (BioRad) and equimolar amounts of cells per condition were labelled with 10X Genomics CellPlex barcodes according to the manufacturer’s protocol (CG000391_Rev B). After pooling, the single cell suspension was further processed using the Chromium Next GEM Single Cell 3’ Reagent Kit v3.1 (CG000388_Rev B) and the Chromium Controller iX (10X Genomics) according to the manufacturer’s protocol.

A total of 40.000 cells (3.200 cells per condition) were used for library preparation. Libraries were sequenced on a NovaSeq X Plus sequencer (Illumina) at approx. 600m reads for the Gene Expression library and 100m reads for the Feature Barcode library.

### Exome Sequencing

RNA was isolated from cultured cells using the Allprep Kit (Qiagen), following manufactureŕs instructions. Exome libraries were produced using SureSelect XT HS2 reagents with UDI-UMI Primers (Agilent). Libraries were sequenced using a NovaSeq X Plus sequencer (Illumina) at mean coverage exceeding 30-fold for parental samples and 200-fold for RASi-resistant samples. The data was mapped using bwa-mem2 version 2.2.1(*61*), on the human genome release 38 complemented with decoys & viral sequences (GRCh38.d1.vd1). The Agilent’s AGeNT software "trimmer" version 3.0.5 & "creak" version 1.0.5 were applied to process molecular barcodes, as suggested by Agilent’s protocols. The base qualities of resulting alignments were then re-scaled using GATK BaseRecalibrator & ApplyBQSR version 4.4.0. The somatic variants were called using GATK’s Mutect2 (version 4.4.0), using the GATK best practice files for panel of normals, germline resource to filter out possible germline variants und common variants for estimating possible contamination. The resulting variants were then annotated with ENSEMBL’s VEP version 102(*62*). An additional filtration step was applied, removing variants with low support (less than 50 reads covering the variant locus, or less than 5 reads supporting the variant), or failing the DKFZ bias filter (github.com/DKFZ-ODCF/DKFZBiasFilter). The copy number alterations were called using cnvkit version 0.9.9 (*63*) with circular binary segmentation (*64, 65*). The calling was done using paired samples, without a panel of normals.

## Acknowledgements

We acknowledge excellent services by the BIH mass and flow cytometry, Bioportal and Genomics core units. This work was partially funded by grants from Deutsche Forschungsgemeinschaft (MO2783/5 to MM, CompCancer graduate school to VH, RAG, NB), and the Einstein Foundation to CJD.

## Data and materials availability

Code, raw data and other data that support the findings of this study are available from the corresponding author upon request, in accordance with national legislation.

## Author contributions

Conceptualisation: MM, CJD, OA; Methodology: OA, MM, EB, VH, RAG; Formal Analysis: OA, EB, VH, RAG, BK; Investigation: OA, ML, AT; Visualisation: OA, EB, VH, RAG; Funding acquisition: MM, CS, NB; Resources: CJD, CS, BP, DH; Project Administration: MM, AT; Supervision: MM, NB, DB; Writing – original draft: MM; Writing – review & editing: all authors

## References

1. W. Kolch, D. Berta, E. Rosta, Dynamic regulation of RAS and RAS signaling. Biochemical Journal 480 (2023), doi:10.1042/BCJ20220234.

2. M. E. Bahar, H. J. Kim, D. R. Kim, Targeting the RAS/RAF/MAPK pathway for cancer therapy: from mechanism to clinical studies. Signal Transduct Target Ther 8 (2023), doi:10.1038/s41392-023-01705-z.

3. H. Lavoie, J. Gagnon, M. Therrien, ERK signalling: a master regulator of cell behaviour, life and fate. Nat Rev Mol Cell Biol 21 (2020), doi:10.1038/s41580-020-0255-7.

4. J. E. Klomp, J. N. Diehl, J. A. Klomp, A. C. Edwards, R. Yang, A. J. Morales, K. E. Taylor, K. Drizyte-Miller, K. L. Bryant, A. Schaefer, J. L. Johnson, E. M. Huntsman, T. M. Yaron, M. Pierobon, E. Baldelli, A. W. Prevatte, N. K. Barker, L. E. Herring, E. F. Petricoin, L. M. Graves, L. C. Cantley, A. D. Cox, C. J. Der, C. A. Stalnecker, Determining the ERK-regulated phosphoproteome driving KRAS-mutant cancer. Science 384, eadk0850 (2024).

5. The Cancer Genome Atlas Network, Comprehensive molecular characterization of human colon and rectal cancer. Nature 487, 330–337 (2012).

6. A. Cervantes, R. Adam, S. Roselló, D. Arnold, N. Normanno, J. Taïeb, J. Seligmann, T. De Baere, P. Osterlund, T. Yoshino, E. Martinelli, Metastatic colorectal cancer: ESMO Clinical Practice Guideline for diagnosis, treatment and follow-up . Annals of Oncology 34 (2023), doi:10.1016/j.annonc.2022.10.003.

7. A. Prahallad, C. Sun, S. Huang, F. Di Nicolantonio, R. Salazar, D. Zecchin, R. L. Beijersbergen, A. Bardelli, R. Bernards, Unresponsiveness of colon cancer to BRAF(V600E) inhibition through feedback activation of EGFR. Nature 483, 100–103 (2012).

8. B. Klinger, A. Sieber, R. Fritsche-Guenther, F. Witzel, L. Berry, D. Schumacher, Y. Yan, P. Durek, M. Merchant, R. Schäfer, C. Sers, N. Blüthgen, Network quantification of EGFR signaling unveils potential for targeted combination therapy. Mol Syst Biol 9, 673 (2013).

9. A. D. Cox, S. W. Fesik, A. C. Kimmelman, J. Luo, C. J. Der, Drugging the undruggable RAS: Mission Possible? Nat Rev Drug Discov 13, 828–851 (2014).

10. J. Canon, K. Rex, A. Y. Saiki, C. Mohr, K. Cooke, D. Bagal, K. Gaida, T. Holt, C. G. Knutson, N. Koppada, B. A. Lanman, J. Werner, A. S. Rapaport, T. San Miguel, R. Ortiz, T. Osgood, J. R. Sun, X. Zhu, J. D. McCarter, L. P. Volak, B. E. Houk, M. G. Fakih, B. H. O’Neil, T. J. Price, G. S. Falchook, J. Desai, J. Kuo, R. Govindan, D. S. Hong, W. Ouyang, H. Henary, T. Arvedson, V. J. Cee, J. R. Lipford, The clinical KRAS(G12C) inhibitor AMG 510 drives anti-tumour immunity. Nature 575 (2019), doi:10.1038/s41586-019-1694-1.

11. J. Hallin, L. D. Engstrom, L. Hargi, A. Calinisan, R. Aranda, D. M. Briere, N. Sudhakar, V. Bowcut, B. R. Baer, J. A. Ballard, M. R. Burkard, J. B. Fell, J. P. Fischer, G. P. Vigers, Y. Xue, S. Gatto, J. Fernandez-Banet, A. Pavlicek, K. Velastagui, R. C. Chao, J. Barton, M. Pierobon, E. Baldelli, E. F. Patricoin, D. P. Cassidy, M. A. Marx, I. I. Rybkin, M. L. Johnson, S. H. Ignatius, P. Lito, K. P. Papadopoulos, P. A. Jänne, P. Olson, J. G. Christensen, The KRASG12C inhibitor MRTX849 provides insight toward therapeutic susceptibility of KRAS-mutant cancers in mouse models and patients. Cancer Discov 10 (2020), doi:10.1158/2159-8290.CD-19-1167.

12. Z. Mao, H. Xiao, P. Shen, Y. Yang, J. Xue, Y. Yang, Y. Shang, L. Zhang, X. Li, Y. Zhang, Y. Du, C. C. Chen, R. T. Guo, Y. Zhang, KRAS(G12D) can be targeted by potent inhibitors via formation of salt bridge. Cell Discov 8 (2022), doi:10.1038/s41421-021-00368-w.

13. F. Lotti, A. M. Jarrar, R. K. Pai, M. Hitomi, J. Lathia, A. Mace, G. A. Gantt, K. Sukhdeo, J. DeVecchio, A. Vasanji, P. Leahy, A. B. Hjelmeland, M. F. Kalady, J. N. Rich, Chemotherapy activates cancer-associated fibroblasts to maintain colorectal cancer-initiating cells by IL-17A. Journal of Experimental Medicine (2013), doi:10.1084/jem.20131195.

14. M. M. Awad, S. Liu, I. I. Rybkin, K. C. Arbour, J. Dilly, V. W. Zhu, M. L. Johnson, R. S. Heist, T. Patil, G. J. Riely, J. O. Jacobson, X. Yang, N. S. Persky, D. E. Root, K. E. Lowder, H. Feng, S. S. Zhang, K. M. Haigis, Y. P. Hung, L. M. Sholl, B. M. Wolpin, J. Wiese, J. Christiansen, J. Lee, A. B. Schrock, L. P. Lim, K. Garg, M. Li, L. D. Engstrom, L. Waters, J. D. Lawson, P. Olson, P. Lito, S.-H. I. Ou, J. G. Christensen, P. A. Jänne, A. J. Aguirre, Acquired Resistance to KRAS G12C Inhibition in Cancer . New England Journal of Medicine 384 (2021), doi:10.1056/nejmoa2105281.

15. N. Tanaka, J. J. Lin, C. Li, M. B. Ryan, J. Zhang, L. A. Kiedrowski, A. G. Michel, M. U. Syed, K. A. Fella, M. Sakhi, I. Baiev, D. Juric, J. F. Gainor, S. J. Klempner, J. K. Lennerz, G. Siravegna, L. Bar-Peled, A. N. Hata, R. S. Heist, R. B. Corcoran, Clinical acquired resistance to krasg12c inhibition through a novel kras switch-ii pocket mutation and polyclonal alterations converging on ras–mapk reactivation. Cancer Discov 11 (2021), doi:10.1158/2159-8290.CD-21-0365.

16. T. Koga, K. Suda, T. Fujino, S. Ohara, A. Hamada, M. Nishino, M. Chiba, M. Shimoji, T. Takemoto, T. Arita, M. Gmachl, M. H. Hofmann, J. Soh, T. Mitsudomi, KRAS Secondary Mutations That Confer Acquired Resistance to KRAS G12C Inhibitors, Sotorasib and Adagrasib, and Overcoming Strategies: Insights From In Vitro Experiments. Journal of Thoracic Oncology 16, 1321–1332 (2021).

17. S. Mukhopadhyay, H. Y. Huang, Z. Lin, M. Ranieri, S. Li, S. Sahu, Y. Liu, Y. Ban, K. Guidry, H. Hu, A. Lopez, F. Sherman, Y. J. Tan, Y. T. Lee, A. P. Armstrong, I. Dolgalev, P. Sahu, T. Zhang, W. Lu, N. S. Gray, J. G. Christensen, T. T. Tang, V. Velcheti, A. Khodadadi-Jamayran, K. K. Wong, B. G. Neel, Genome-Wide CRISPR Screens Identify Multiple Synthetic Lethal Targets That Enhance KRASG12C Inhibitor Efficacy. Cancer Res 183 (2023), doi:10.1158/0008-5472.CAN-23-2729.

18. A. C. Edwards, C. A. Stalnecker, A. J. Morales, K. E. Taylor, J. E. Klomp, J. A. Klomp, A. M. Waters, N. Sudhakar, J. Hallin, T. T. Tang, P. Olson, L. Post, J. G. Christensen, A. D. Cox, C. J. Der, TEAD Inhibition Overcomes YAP1/TAZ-Driven Primary and Acquired Resistance to KRASG12C Inhibitors. Cancer Res 183 (2023), doi:10.1158/0008-5472.CAN-23-2994.

19. C. J. Schulze, K. J. Seamon, Y. Zhao, Y. C. Yang, J. Cregg, D. Kim, A. Tomlinson, T. J. Choy, Z. Wang, B. Sang, Y. Pourfarjam, J. Lucas, A. Cuevas-Navarro, C. Ayala-Santos, A. Vides, C. Li, A. Marquez, M. Zhong, V. Vemulapalli, C. Weller, A. Gould, D. M. Whalen, A. Salvador, A. Milin, M. Saldajeno-Concar, N. Dinglasan, A. Chen, J. Evans, J. E. Knox, E. S. Koltun, M. Singh, R. Nichols, D. Wildes, A. L. Gill, J. A. M. Smith, P. Lito, Chemical remodeling of a cellular chaperone to target the active state of mutant KRAS. Science (1979) 381 (2023), doi:10.1126/science.adg9652.

20. M. Holderfield, B. J. Lee, J. Jiang, A. Tomlinson, K. J. Seamon, A. Mira, E. Patrucco, G. Goodhart, J. Dilly, Y. Gindin, N. Dinglasan, Y. Wang, L. P. Lai, S. Cai, L. Jiang, N. Nasholm, N. Shifrin, C. Blaj, H. Shah, J. W. Evans, N. Montazer, O. Lai, J. Shi, E. Ahler, E. Quintana, S. Chang, A. Salvador, A. Marquez, J. Cregg, Y. Liu, A. Milin, A. Chen, T. B. Ziv, D. Parsons, J. E. Knox, J. E. Klomp, J. Roth, M. Rees, M. Ronan, A. Cuevas-Navarro, F. Hu, P. Lito, D. Santamaria, A. J. Aguirre, A. M. Waters, C. J. Der, C. Ambrogio, Z. Wang, A. L. Gill, E. S. Koltun, J. A. M. Smith, D. Wildes, M. Singh, Concurrent inhibition of oncogenic and wild-type RAS-GTP for cancer therapy. Nature (2024), doi:10.1038/s41586-024-07205-6.

21. U. N. Wasko, J. Jiang, A. Curiel-Garcia, Y. Wang, B. Lee, M. Orlen, K. Drizyte-Miller, M. Menard, J. Dilly, S. A. Sastra, C. F. Palermo, T. Dalton, M. C. Hasselluhn, A. R. Decker-Farrell, S. Chang, L. Jiang, X. Wei, Y. C. Yang, C. Helland, H. Courtney, Y. Gindin, R. Zhao, S. B. Kemp, C. Clendenin, R. Sor, W. Vostrejs, A. A. Amparo, P. S. Hibshman, M. G. Rees, M. M. Ronan, J. A. Roth, B. Bakir, M. A. Badgley, J. A. Chabot, M. D. Kluger, G. A. Manji, E. Quintana, Z. Wang, J. A. M. Smith, M. Holderfield, D. Wildes, A. J. Aguirre, C. J. Der, R. H. Vonderheide, B. Z. Stanger, M. Singh, K. P. Olive, Tumor-selective effects of active RAS inhibition in pancreatic ductal adenocarcinoma. bioRxiv (2023).

22. E. J. Morris, S. Jha, C. R. Restaino, P. Dayananth, H. Zhu, A. Cooper, D. Carr, Y. Deng, W. Jin, S. Black, B. Long, J. Liu, E. DiNunzio, W. Windsor, R. Zhang, S. Zhao, M. H. Angagaw, E. M. Pinheiro, J. Desai, L. Xiao, G. Shipps, A. Hruza, J. Wang, J. Kelly, S. Paliwal, X. Gao, B. S. Babu, L. Zhu, P. Daublain, L. Zhang, B. A. Lutterbach, M. R. Pelletier, U. Philippar, P. Siliphaivanh, D. Witter, P. Kirschmeier, W. Robert Bishop, D. Hicklin, D. Gary Gillil, L. Jayaraman, L. Zawel, S. Fawell, A. A. Samatar, Discovery of a novel ERK inhibitor with activity in models of acquired resistance to BRAF and MEK inhibitors. Cancer Discov 3, 742–750 (2013).

23. J. G. Albeck, G. B. Mills, J. S. Brugge, Frequency-Modulated Pulses of ERK Activity Transmit Quantitative Proliferation Signals. Mol Cell 49, 249–261 (2013).

24. J. G. Albeck, G. B. Mills, J. S. Brugge, Frequency-Modulated Pulses of ERK Activity Transmit Quantitative Proliferation Signals. Mol Cell 49, 249–261 (2013).

25. M. L. Samuels, M. J. Weber, J. M. Bishop, M. McMahon, Conditional transformation of cells and rapid activation of the mitogen-activated protein kinase cascade by an estradiol-dependent human raf-1 protein kinase. Mol Cell Biol 13 (1993), doi:10.1128/mcb.13.10.6241.

26. S. P. Suehnholz, M. H. Nissan, H. Zhang, R. Kundra, S. Nandakumar, C. Lu, S. Carrero, A. Dhaneshwar, N. Fernandez, B. W. Xu, M. E. Arcila, A. Zehir, A. Syed, A. R. Brannon, J. E. Rudolph, E. Paraiso, P. J. Sabbatini, R. L. Levine, A. Dogan, J. Gao, M. Ladanyi, A. Drilon, M. F. Berger, D. B. Solit, N. Schultz, D. Chakravarty, Quantifying the Expanding Landscape of Clinical Actionability for Patients with Cancer. Cancer Discov 14 (2024), doi:10.1158/2159-8290.CD-23-0467.

27. I. C. Cirstea, L. Gremer, R. Dvorsky, S. C. Zhang, R. P. Piekorz, M. Zenker, M. R. Ahmadian, Diverging gain-of-function mechanisms of two novel KRAS mutations associated with Noonan and cardio-facio-cutaneous syndromes. Hum Mol Genet 22 (2013), doi:10.1093/hmg/dds426.

28. T. Kobayashi, Y. Aoki, T. Niihori, H. Cavé, A. Verloes, N. Okamoto, H. Kawame, I. Fujiwara, F. Takada, T. Ohata, S. Sakazume, T. Ando, N. Nakagawa, P. Lapunzina, A. G. Meneses, G. Gillessen-Kaesbach, D. Wieczorek, K. Kurosawa, S. Mizuno, H. Ohashi, A. David, N. Philip, A. Guliyeva, Y. Narumi, S. Kure, S. Tsuchiya, Y. Matsubara, Molecular and clinical analysis of RAF1 in Noonan syndrome and related disorders: Dephosphorylation of serine 259 as the essential mechanism for mutant activation. Hum Mutat 31 (2010), doi:10.1002/humu.21187.

29. M. Holderfield, B. J. Lee, J. Jiang, A. Tomlinson, K. J. Seamon, A. Mira, E. Patrucco, G. Goodhart, J. Dilly, Y. Gindin, N. Dinglasan, Y. Wang, L. P. Lai, S. Cai, L. Jiang, N. Nasholm, N. Shifrin, C. Blaj, H. Shah, J. W. Evans, N. Montazer, O. Lai, J. Shi, E. Ahler, E. Quintana, S. Chang, A. Salvador, A. Marquez, J. Cregg, Y. Liu, A. Milin, A. Chen, T. B. Ziv, D. Parsons, J. E. Knox, J. E. Klomp, J. Roth, M. Rees, M. Ronan, A. Cuevas-Navarro, F. Hu, P. Lito, D. Santamaria, A. J. Aguirre, A. M. Waters, C. J. Der, C. Ambrogio, Z. Wang, A. L. Gill, E. S. Koltun, J. A. M. Smith, D. Wildes, M. Singh, Concurrent inhibition of oncogenic and wild-type RAS-GTP for cancer therapy. Nature (2024), doi:10.1038/s41586-024-07205-6.

30. S. Nakhaei-Rad, F. Haghighi, F. Bazgir, J. Dahlmann, A. V. Busley, M. Buchholzer, K. Kleemann, A. Schänzer, A. Borchardt, A. Hahn, S. Kötter, D. Schanze, R. Anand, F. Funk, A. V. Kronenbitter, J. Scheller, R. P. Piekorz, A. S. Reichert, M. Volleth, M. J. Wolf, I. C. Cirstea, B. D. Gelb, M. Tartaglia, J. P. Schmitt, M. Krüger, I. Kutschka, L. Cyganek, M. Zenker, G. Kensah, M. R. Ahmadian, Molecular and cellular evidence for the impact of a hypertrophic cardiomyopathy-associated RAF1 variant on the structure and function of contractile machinery in bioartificial cardiac tissues. Commun Biol 6 (2023), doi:10.1038/s42003-023-05013-8.

31. K. Kondoh, K. Terasawa, H. Morimoto, E. Nishida, Regulation of Nuclear Translocation of Extracellular Signal-Regulated Kinase 5 by Active Nuclear Import and Export Mechanisms. Mol Cell Biol 26 (2006), doi:10.1128/mcb.26.5.1679-1690.2006.

32. A. J. King, H. Sun, B. Diaz, D. Barnard, W. Miao, S. Bagrodia, M. S. Marshall, The protein kinase Pak3 positively regulates Raf-1 activity through phosphorylation of serine 338. Nature 396 (1998), doi:10.1038/24184.

33. A. Gregorieff, Y. Liu, M. R. Inanlou, Y. Khomchuk, J. L. Wrana, Yap-dependent reprogramming of Lgr5+ stem cells drives intestinal regeneration and cancer. Nature (2015), doi:10.1038/nature15382.

34. Y. Wang, X. Xu, D. Maglic, M. T. Dill, K. Mojumdar, P. K. S. Ng, K. J. Jeong, Y. H. Tsang, D. Moreno, V. H. Bhavana, X. Peng, Z. Ge, H. Chen, J. Li, Z. Chen, H. Zhang, L. Han, D. Du, C. J. Creighton, G. B. Mills, The Cancer Genome Atlas Research Network, F. Camargo, H. Liang, Comprehensive Molecular Characterization of the Hippo Signaling Pathway in Cancer. Cell Rep 25 (2018), doi:10.1016/j.celrep.2018.10.001.

35. D. Serra, U. Mayr, A. Boni, I. Lukonin, M. Rempfler, L. Challet Meylan, M. B. Stadler, P. Strnad, P. Papasaikas, D. Vischi, A. Waldt, G. Roma, P. Liberali, Self-organization and symmetry breaking in intestinal organoid development. Nature 569, 66–72 (2019).

36. A. Merlos-Suárez, F. M. Barriga, P. Jung, M. Iglesias, M. V. Céspedes, D. Rossell, M. Sevillano, X. Hernando-Momblona, V. da Silva-Diz, P. Muñoz, H. Clevers, E. Sancho, R. Mangues, E. Batlle, The intestinal stem cell signature identifies colorectal cancer stem cells and predicts disease relapse. Cell Stem Cell 8, 511–524 (2011).

37. A. Álvarez-Varela, L. Novellasdemunt, F. M. Barriga, X. Hernando-Momblona, A. Cañellas-Socias, S. Cano-Crespo, M. Sevillano, C. Cortina, D. Stork, C. Morral, G. Turon, F. Slebe, L. Jiménez-Gracia, G. Caratù, P. Jung, G. Stassi, H. Heyn, D. V. F. Tauriello, L. Mateo, S. Tejpar, E. Sancho, C. Stephan-Otto Attolini, E. Batlle, Mex3a marks drug-tolerant persister colorectal cancer cells that mediate relapse after chemotherapy. Nat Cancer 3 (2022), doi:10.1038/s43018-022-00402-0.

38. A. Liberzon, C. Birger, H. Thorvaldsdóttir, M. Ghandi, J. P. Mesirov, P. Tamayo, The Molecular Signatures Database Hallmark Gene Set Collection. Cell Syst 1, 417–425 (2015).

39. S. Li, A. Balmain, C. M. Counter, A model for RAS mutation patterns in cancers: finding the sweet spot Nat Rev Cancer 18 (2018), doi:10.1038/s41568-018-0076-6.

40. R. Yaeger, R. Mezzadra, J. Sinopoli, Y. Bian, M. Marasco, E. Kaplun, Y. Gao, H. Zhao, A. D. C. Paula, Y. Zhu, A. C. Perez, K. Chadalavada, E. Tse, S. Chowdhry, S. Bowker, Q. Chang, B. Qeriqi, B. Weigelt, G. J. Nanjangud, M. F. Berger, H. Der-Torossian, K. Anderes, N. D. Socci, J. Shia, G. J. Riely, Y. R. Murciano-Goroff, B. T. Li, J. G. Christensen, J. S. Reis-Filho, D. B. Solit, E. de Stanchina, S. W. Lowe, N. Rosen, S. Misale, Molecular Characterization of Acquired Resistance to KRASG12C–EGFR Inhibition in Colorectal Cancer. Cancer Discov 13 (2023), doi:10.1158/2159-8290.CD-22-0405.

41. I. El Bouchikhi, K. Belhassan, F. Z. Moufid, M. Iraqui Houssaini, L. Bouguenouch, I. Samri, S. Atmani, K. Ouldim, Noonan syndrome-causing genes: Molecular update and an assessment of the mutation rate. Int J Pediatr Adolesc Med 3 (2016), doi:10.1016/j.ijpam.2016.06.003.

42. B. Pandit, A. Sarkozy, L. A. Pennacchio, C. Carta, K. Oishi, S. Martinelli, E. A. Pogna, W. Schackwitz, A. Ustaszewska, A. Landstrom, J. M. Bos, S. R. Ommen, G. Esposito, F. Lepri, C. Faul, P. Mundel, J. P. López Siguero, R. Tenconi, A. Selicorni, C. Rossi, L. Mazzanti, I. Torrente, B. Marino, M. C. Digilio, G. Zampino, M. J. Ackerman, B. Dallapiccola, M. Tartaglia, B. D. Gelb, Gain-of-function RAF1 mutations cause Noonan and LEOPARD syndromes with hypertrophic cardiomyopathy. Nat Genet 39, 1007– 1012 (2007).

43. M. Molzan, B. Schumacher, C. Ottmann, A. Baljuls, L. Polzien, M. Weyand, P. Thiel, R. Rose, M. Rose, P. Kuhenne, M. Kaiser, U. R. Rapp, J. Kuhlmann, C. Ottmann, Impaired Binding of 14-3-3 to C- RAF in Noonan Syndrome Suggests New Approaches in Diseases with Increased Ras Signaling. Mol Cell Biol 30 (2010), doi:10.1128/mcb.01636-09.

44. J. E. Klomp, J. A. Klomp, C. J. Der, The ERK mitogen-activated protein kinase signaling network: The final frontier in RAS signal transduction Biochem Soc Trans 49 (2021), doi:10.1042/BST20200507.

45. J. A. Klomp, J. E. Klomp, C. A. Stalnecker, K. L. Bryant, A. C. Edwards, K. Drizyte-Miller, P. S. Hibshman, J. N. Diehl, Y. S. Lee, A. J. Morales, K. E. Taylor, S. Peng, N. L. Tran, L. E. Herring, A. W. Prevatte, N. K. Barker, L. D. Hover, J. Hallin, S. Chowdhury, O. Coker, H. M. Lee, C. M. Goodwin, P. Gautam, P. Olson, J. G. Christensen, J. P. Shen, S. Kopetz, L. M. Graves, K. H. Lim, A. Wang-Gillam, K. Wennerberg, A. D. Cox, C. J. Der, Defining the KRAS- and ERK-dependent transcriptome in KRAS-mutant cancers. Science 384, eadk0775 (2024).

46. A. Ram, D. Murphy, N. DeCuzzi, M. Patankar, J. Hu, M. Pargett, J. G. Albeck, A guide to ERK dynamics, part 1: mechanisms and models. Biochemical Journal 480, 1887–1907 (2023).

47. A. Ram, D. Murphy, N. DeCuzzi, M. Patankar, J. Hu, M. Pargett, J. G. Albeck, A guide to ERK dynamics, part 2: downstream decoding*Biochemical* Journal 148 (2023), doi:10.1042/BCJ20230277.

48. G. Bollag, P. Hirth, J. Tsai, J. Zhang, P. N. Ibrahim, H. Cho, W. Spevak, C. Zhang, Y. Zhang, G. Habets, E. A. Burton, B. Wong, G. Tsang, B. L. West, B. Powell, R. Shellooe, A. Marimuthu, H. Nguyen, K. Y. J. Zhang, D. R. Artis, J. Schlessinger, F. Su, B. Higgins, R. Iyer, K. Dandrea, A. Koehler, M. Stumm, P. S. Lin, R. J. Lee, J. Grippo, I. Puzanov, K. B. Kim, A. Ribas, G. A. McArthur, J. A. Sosman, P. B. Chapman, K. T. Flaherty, X. Xu, K. L. Nathanson, K. Nolop, Clinical efficacy of a RAF inhibitor needs broad target blockade in BRAF-mutant melanoma. Nature (2010), doi:10.1038/nature09454.

49. S. Cagnol, N. Rivard, Oncogenic KRAS and BRAF activation of the MEK/ERK signaling pathway promotes expression of dual-specificity phosphatase 4 (DUSP4/MKP2) resulting in nuclear ERK1/2 inhibition. Oncogene 32, 564–576 (2013).

50. Y. Liang, F. Sheikh, Scaffold proteins regulating extracellular regulated kinase function in cardiac hypertrophy and disease Front Pharmacol 7 (2016), doi:10.3389/fphar.2016.00037.

51. A. Martín-Vega, L. Ruiz-Peinado, R. García-Gómez, A. Herrero, D. de la Fuente-Vivas, S. Parvathaneni, R. Caloto, M. Morante, A. von Kriegsheim, X. R. Bustelo, D. B. Sacks, B. Casar, P. Crespo, Scaffold coupling: ERK activation by transphosphorylation across different scaffold protein species. Sci Adv 9 (2023), doi:10.1126/sciadv.add7969.

52. M. Adachi, M. Fukuda, E. Nishida, Nuclear export of MAP kinase (ERK) involves a MAP kinase kinase (MEK)-dependent active transport mechanism. Journal of Cell Biology 148 (2000), doi:10.1083/jcb.148.5.849.

53. E. Formstecher, J. W. Ramos, M. Fauquet, D. A. Calderwood, J. C. Hsieh, B. Canton, X. T. Nguyen, J. V. Barnier, J. Camonis, M. H. Ginsberg, H. Chneiweiss, PEA-15 Mediates Cytoplasmic Sequestration of ERK MAP Kinase. Dev Cell 1 (2001), doi:10.1016/S1534-5807(01)00035-1.

54. A. W. Whitehurst, J. L. Wilsbacher, Y. You, K. Luby-Phelps, M. S. Moore, M. H. Cobb, ERK2 enters the nucleus by a carrier-independent mechanism. Proc Natl Acad Sci U S A 99 (2002), doi:10.1073/pnas.112495999.

55. W. Brugger, M. Thomas, EGFR-TKI resistant non-small cell lung cancer (NSCLC): New developments and implications for future treatment. Lung Cancer 77, 2–8 (2012).

56. I. Ozkan-Dagliyan, J. N. Diehl, S. D. George, A. Schaefer, B. Papke, K. Klotz-Noack, A. M. Waters, C. M. Goodwin, P. Gautam, M. Pierobon, S. Peng, T. S. K. Gilbert, K. H. Lin, O. Dagliyan, K. Wennerberg, E. F. Petricoin, N. L. Tran, S. V. Bhagwat, R. V. Tiu, S. Bin Peng, L. E. Herring, L. M. Graves, C. Sers, K. C. Wood, A. D. Cox, C. J. Der, Low-Dose Vertical Inhibition of the RAF-MEK-ERK Cascade Causes Apoptotic Death of KRAS Mutant Cancers. Cell Rep (2020), doi:10.1016/j.celrep.2020.107764.

57. S. Zheng, W. Wang, J. Aldahdooh, A. Malyutina, T. Shadbahr, Z. Tanoli, A. Pessia, J. Tang, SynergyFinder Plus: Toward Better Interpretation and Annotation of Drug Combination Screening Datasets. Genomics Proteomics Bioinformatics 20 (2022), doi:10.1016/j.gpb.2022.01.004.

58. R. Brandt, T. Sell, M. Lüthen, F. Uhlitz, B. Klinger, P. Riemer, C. Giesecke-Thiel, S. Schulze, I. A. El-Shimy, D. Kunkel, B. Fauler, T. Mielke, N. Mages, B. G. Herrmann, C. Sers, N. Blüthgen, M. Morkel, Cell type-dependent differential activation of ERK by oncogenic KRAS in colon cancer and intestinal epithelium. Nat Commun 10, 2919 (2019).

59. R. Astaburuaga-García, T. Sell, S. Mutlu, A. Sieber, K. Lauber, N. Blüthgen, RUCova: Removal of Unwanted Covariance in mass cytometry data. bioRxiv, 2024.05.24.595717 (2024).

60. M. E. Ritchie, B. Phipson, D. Wu, Y. Hu, C. W. Law, W. Shi, G. K. Smyth, Limma powers differential expression analyses for RNA-sequencing and microarray studies. Nucleic Acids Res 43 (2015), doi:10.1093/nar/gkv007.

61. V. Md, S. Misra, H. Li, S. Aluru, in *Proceedings - 2019 IEEE 33rd International Parallel and Distributed Processing Symposium*, IPDPS 2019, (2019).

62. W. McLaren, L. Gil, S. E. Hunt, H. S. Riat, G. R. S. Ritchie, A. Thormann, P. Flicek, F. Cunningham, The Ensembl Variant Effect Predictor. Genome Biol 17 (2016), doi:10.1186/s13059-016-0974-4.

63. E. Talevich, A. H. Shain, T. Botton, B. C. Bastian, CNVkit: Genome-Wide Copy Number Detection and Visualization from Targeted DNA Sequencing. PLoS Comput Biol 12 (2016), doi:10.1371/journal.pcbi.1004873.

64. A. B. Olshen, H. Bengtsson, P. Neuvial, P. T. Spellman, R. A. Olshen, V. E. Seshan, Parent-specific copy number in paired tumor-normal studies using circular binary segmentation. Bioinformatics 27 (2011), doi:10.1093/bioinformatics/btr329.

65. E. S. Venkatraman, A. B. Olshen, A faster circular binary segmentation algorithm for the analysis of array CGH data. Bioinformatics 23 (2007), doi:10.1093/bioinformatics/btl646.

