## Supplemental Figures 1-6 for "Reporter-based screening identifies RAS-RAF mutations as drivers of resistance to active-state RAS inhibition in colorectal cancer"

**Supplementary Figures**

Authors: Oleksandra Aust[1], Eric Blanc[2], Mareen Lüthen[1], Viola Hollek[1], Rosario Astaburuaga-García[1], Bertram Klinger[1], Alexandra Trinks[3], Dieter Beule[2], Björn Papke[1, 4, 5], David Horst[1, 4], Nils Blüthgen[1, 4], Christine Sers[1, 4], Channing J. Der[5], Markus Morkel[1, 3, 4]

Affiliations:

1 Institute of Pathology, Charité - Universitätsmedizin Berlin, Corporate Member of Freie Universität Berlin and Humboldt-Universität zu Berlin, Charitéplatz 1, 10117, Berlin, Germany

2 Core Unit Bioinformatics (CUBI), Berlin Institute of Health at Charité - Universitätsmedizin Berlin, 10117 Berlin, Germany

3 Bioportal Single Cells, Berlin Institute of Health at Charité - Universitätsmedizin Berlin, 10117 Berlin, Germany.

4 German Cancer Consortium (DKTK) Partner Site Berlin, German Cancer Research Center (DKFZ), 69120 Heidelberg, Germany

5 University of North Carolina at Chapel Hill, Chapel Hill, North Carolina, USA

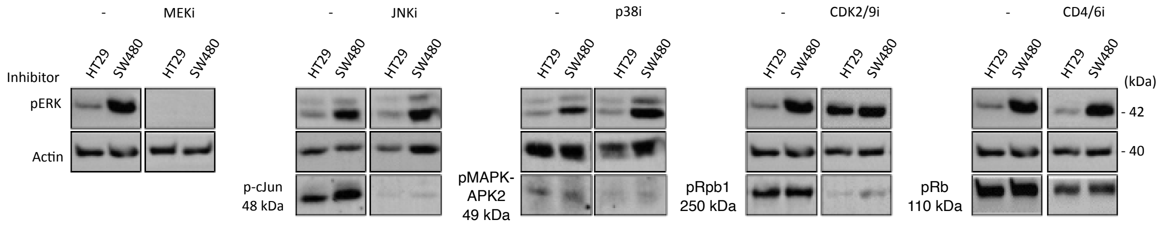


**Supplementary Figure 1: Reporter specificity tests, using Western blot analysis for ERK and established targets of the respective inhibitors**. Cell lines used and inhibitors are given. For details, see Materials and Methods. Insets per inhibitor are from same membrane.

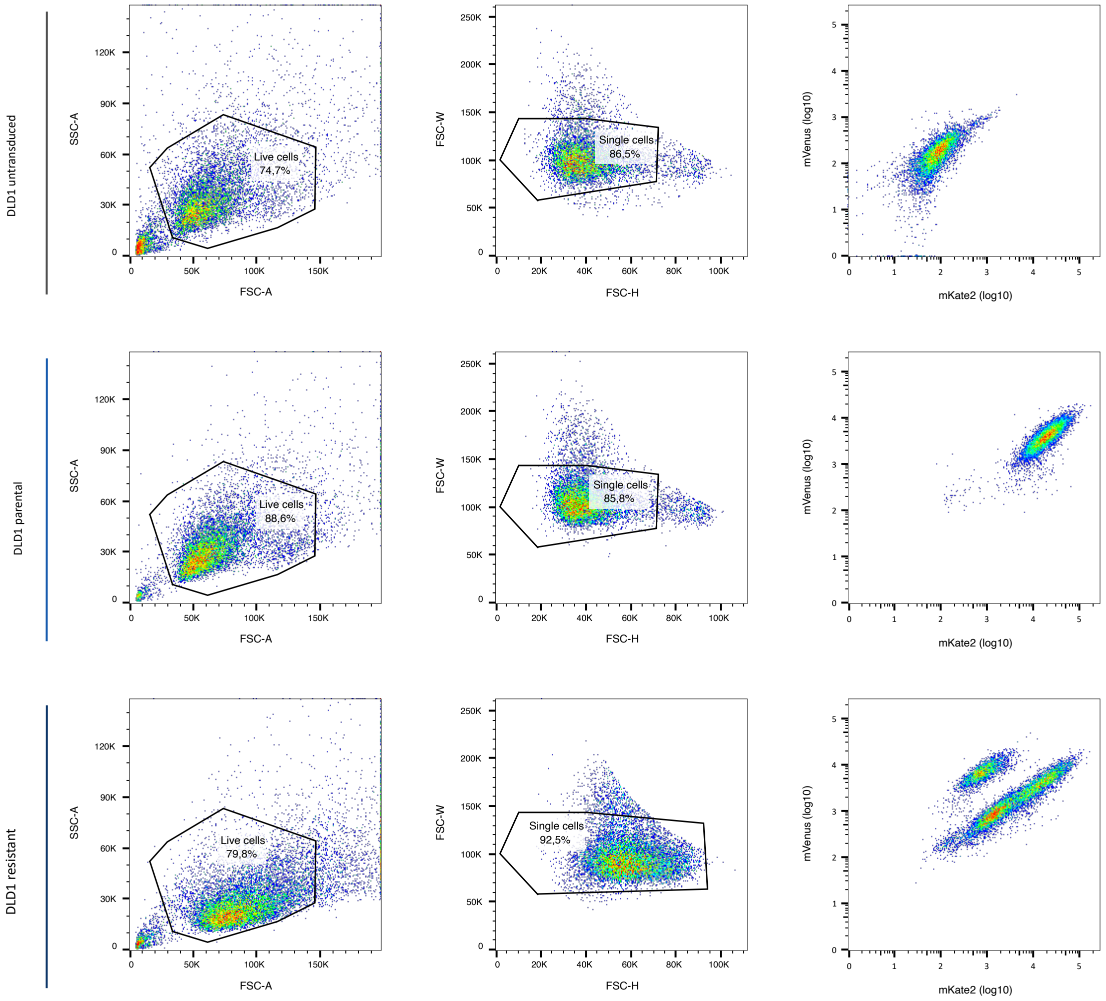


**Supplementary Figure 2: Gating strategy for sorting of RASi-resistant DLD1 subpopulations**. **Top to bottom**: Untransfected DLD1 cells; parental DLD1 cell population transfected with the FIREX plasmids; resistant DLD1 population after dose escalation with RMC-7977. **Left to right:** All sorting events with sorting window for Live cells; All live cells with sorting window for single cells; mVenus/mKate2 fluorescence of cells in the single cells window. For sorting windows, see main Fig. 3H.

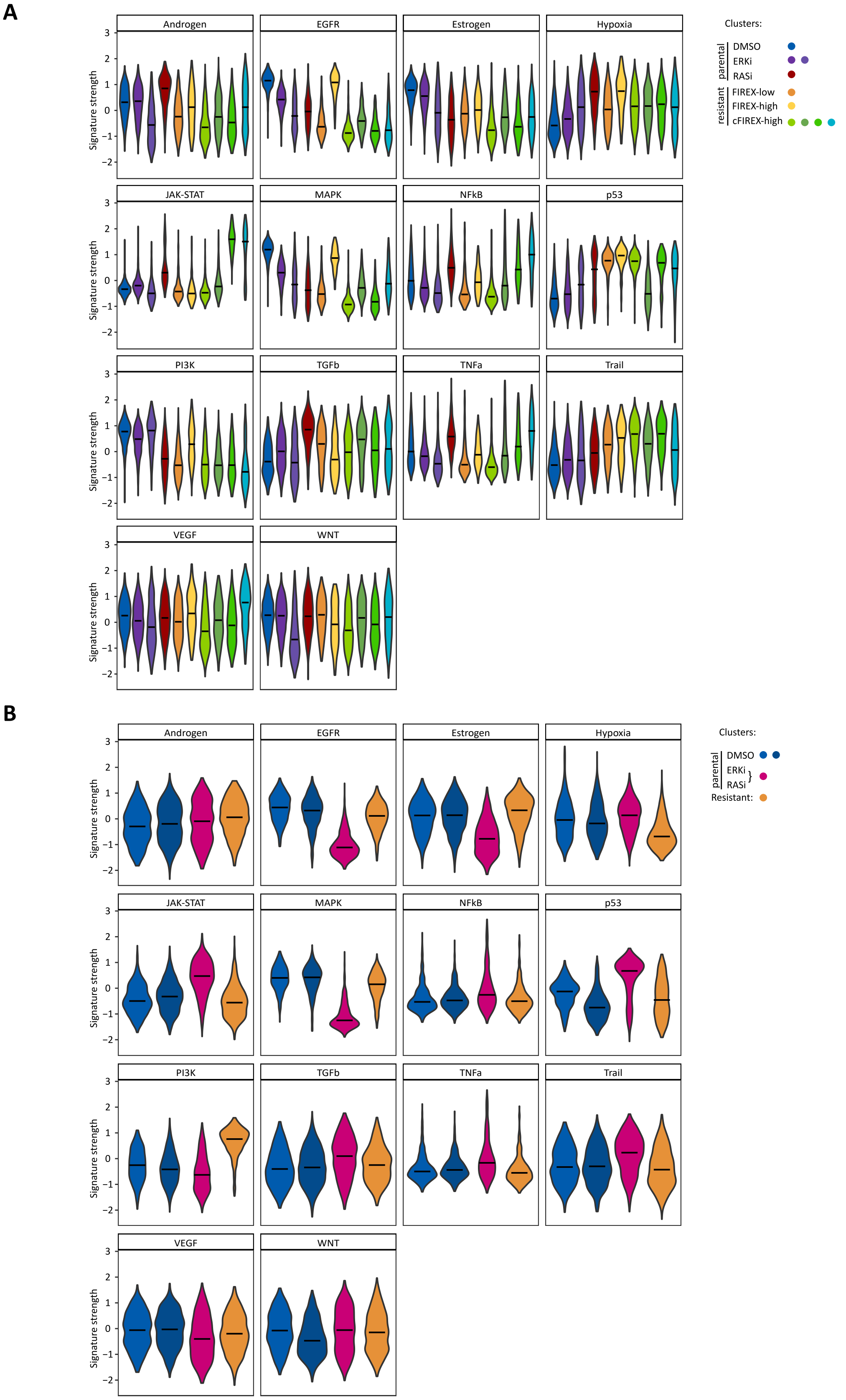


**Supplementary Figure 3**: **Complete Progeny pathway activity analysis of clusters, as shown in main Figure 5D A** Pathway scores *(1)* for clusters of DLD1 cells, **B** Pathway scores for clusters of SW480 cells.

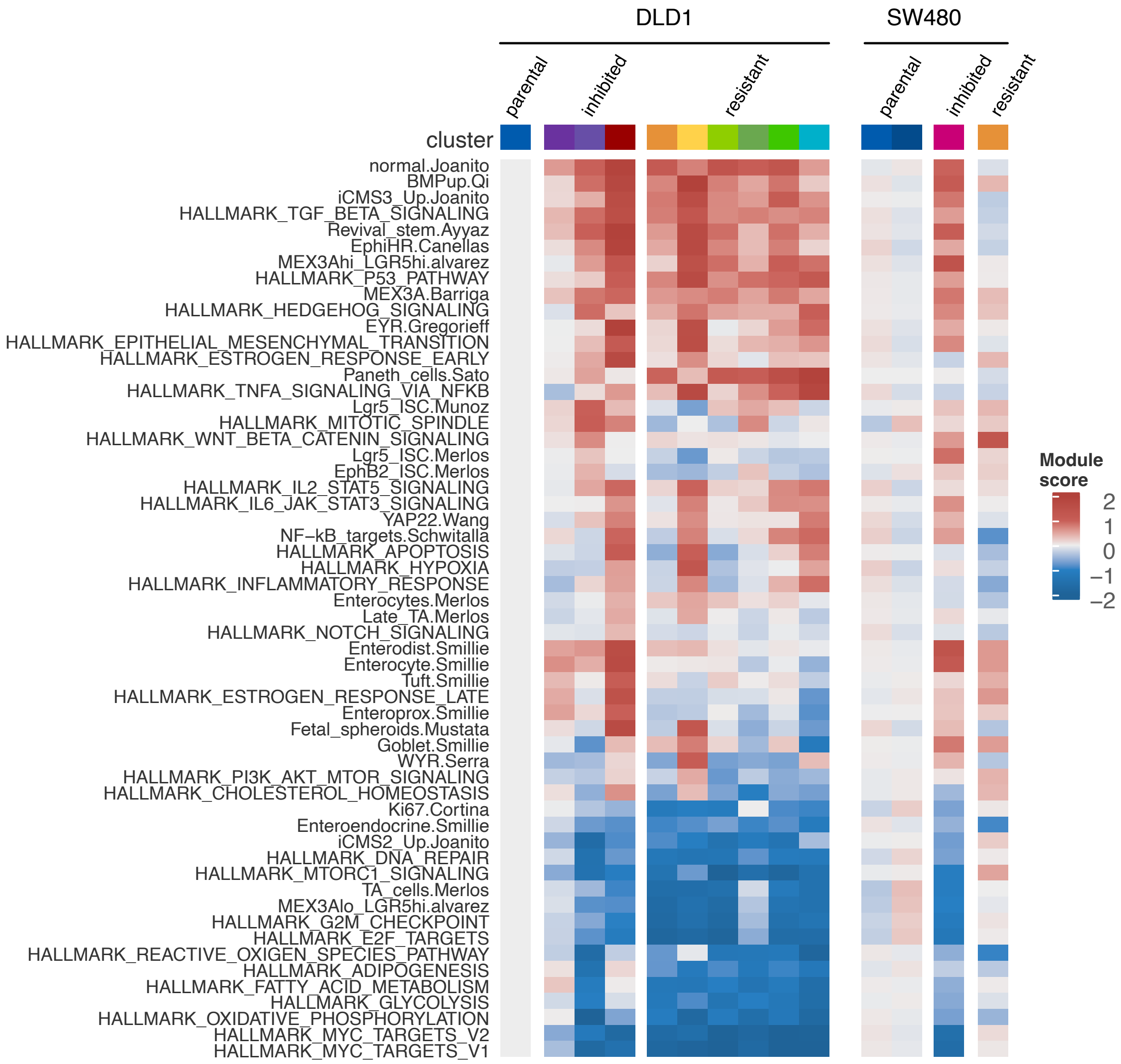


**Supplementary Figure 4: Activities of CRC-relevant signaling activities.** Figure gives mean pathway score per signature. For details on signatures, see associated Information on Zenodo. Signatures were extracted from the following papers: For hallmark signatures *(2)*, for Progeny, for signatures related to colon and CRC cell heterogeneity *(3–17).*

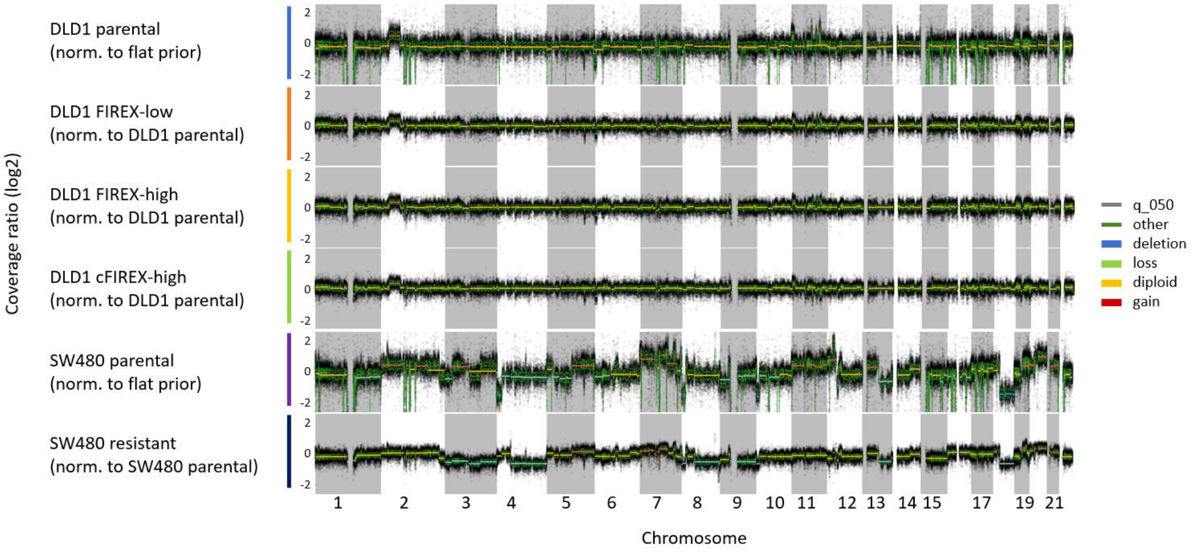


**Supplementary Figure 5: Copy number profiles of parental and resistant cell subpopulations, as indicated.** For gene lists with copy number changes, refer to Supplementary table 1. Parental lines were compared to a flat (diploid) prior, while resistant cell lines were normalized to the parental line.

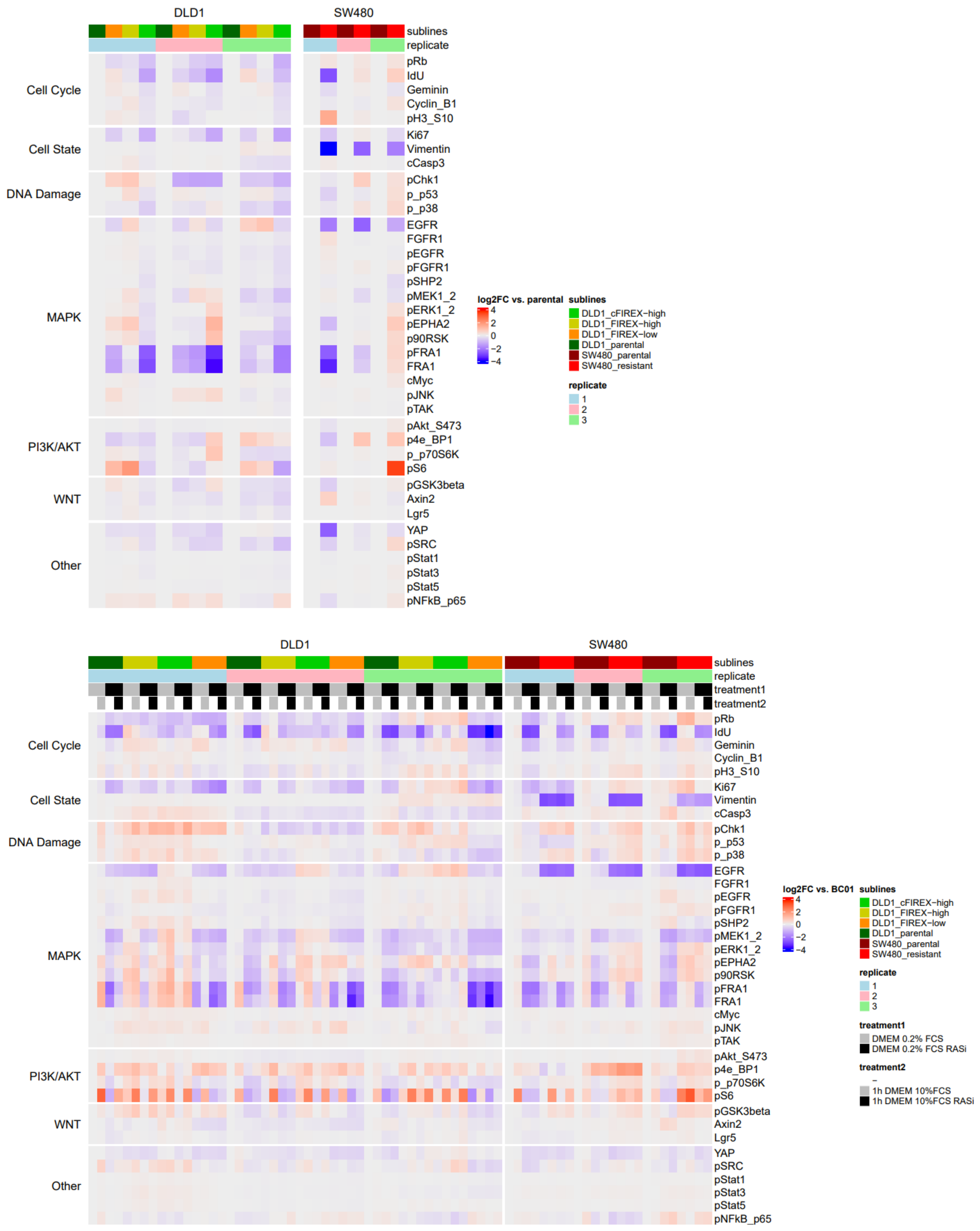


**Supplementary Figure 6: Information on mass cytometry replicate experiments. Above:** Biological replicate mass cytometry experiments without perturbation (relates to main Figure 6A), **Below:** Biological replicate mass cytometry experiments with perturbation (relates to main Figure 6C).

**Bibliography for Supplementary Figures:**


1. M. Schubert, B. Klinger, M. Klünemann, A. Sieber, F. Uhlitz, S. Sauer, M. J. Garnett, N. Blüthgen, J. Saez-Rodriguez, Perturbation-response genes reveal signaling footprints in cancer gene expression. *Nat Commun* **9**, 20 (2018).

2. A. Subramanian, P. Tamayo, V. K. Mootha, S. Mukherjee, B. L. Ebert, M. A. Gillette, A. Paulovich, S. L. Pomeroy, T. R. Golub, E. S. Lander, J. P. Mesirov, Gene set enrichment analysis: a knowledge-based approach for interpreting genome-wide expression profiles. *Proc Natl Acad Sci U S A* **102**, 15545–15550 (2005).

3. S. Schwitalla, A. A. Fingerle, P. Cammareri, T. Nebelsiek, S. I. Göktuna, P. K. Ziegler, O. Canli, J. Heijmans, D. J. Huels, G. Moreaux, R. A. Rupec, M. Gerhard, R. Schmid, N. Barker, H. Clevers, R. Lang, J. Neumann, T. Kirchner, M. M. Taketo, G. R. van den Brink, O. J. Sansom, M. C. Arkan, F. R. Greten, Intestinal Tumorigenesis Initiated by Dedifferentiation and Acquisition of Stem-Cell-like Properties. *Cell* **152**, 25–38 (2013).

4. R. C. Mustata, G. Vasile, V. Fernandez-Vallone, S. Strollo, A. Lefort, F. Libert, D. Monteyne, D. Pérez-Morga, G. Vassart, M. I. Garcia, Identification of Lgr5-Independent Spheroid-Generating Progenitors of the Mouse Fetal Intestinal Epithelium. *Cell Rep* **5** (2013), doi:10.1016/j.celrep.2013.09.005.

5. A. Álvarez-Varela, L. Novellasdemunt, F. M. Barriga, X. Hernando-Momblona, A. Cañellas-Socias, S. Cano-Crespo, M. Sevillano, C. Cortina, D. Stork, C. Morral, G. Turon, F. Slebe, L. Jiménez-Gracia, G. Caratù, P. Jung, G. Stassi, H. Heyn, D. V. F. Tauriello, L. Mateo, S. Tejpar, E. Sancho, C. Stephan-Otto Attolini, E. Batlle, Mex3a marks drug-tolerant persister colorectal cancer cells that mediate relapse after chemotherapy. *Nat Cancer* **3** (2022), doi:10.1038/s43018-022-00402-0.

6. A. Gregorieff, Y. Liu, M. R. Inanlou, Y. Khomchuk, J. L. Wrana, Yap-dependent reprogramming of Lgr5+ stem cells drives intestinal regeneration and cancer. *Nature* (2015), doi:10.1038/nature15382.

7. Y. Wang, X. Xu, D. Maglic, M. T. Dill, K. Mojumdar, P. K. S. Ng, K. J. Jeong, Y. H. Tsang, D. Moreno, V. H. Bhavana, X. Peng, Z. Ge, H. Chen, J. Li, Z. Chen, H. Zhang, L. Han, D. Du, C. J. Creighton, G. B. Mills, The Cancer Genome Atlas Research Network, F. Camargo, H. Liang, Comprehensive Molecular Characterization of the Hippo Signaling Pathway in Cancer. *Cell Rep* **25** (2018), doi:10.1016/j.celrep.2018.10.001.

8. D. Serra, U. Mayr, A. Boni, I. Lukonin, M. Rempfler, L. Challet Meylan, M. B. Stadler, P. Strnad, P. Papasaikas, D. Vischi, A. Waldt, G. Roma, P. Liberali, Self-organization and symmetry breaking in intestinal organoid development. *Nature* **569**, 66–72 (2019).

9. T. Sato, J. H. van Es, H. J. Snippert, D. E. Stange, R. G. Vries, M. van den Born, N. Barker, N. F. Shroyer, M. van de Wetering, H. Clevers, Paneth cells constitute the niche for Lgr5 stem cells in intestinal crypts. *Nature* **469**, 415–418 (2010).

10. A. Merlos-Suárez, F. M. Barriga, P. Jung, M. Iglesias, M. V. Céspedes, D. Rossell, M. Sevillano, X. Hernando-Momblona, V. da Silva-Diz, P. Muñoz, H. Clevers, E. Sancho, R. Mangues, E. Batlle, The intestinal stem cell signature identifies colorectal cancer stem cells and predicts disease relapse. *Cell Stem Cell* **8**, 511–524 (2011).

11. F. M. Barriga, E. Montagni, M. Mana, M. Mendez-Lago, X. Hernando-Momblona, M. Sevillano, A. Guillaumet-Adkins, G. Rodriguez-Esteban, S. J. A. Buczacki, M. Gut, H. Heyn, D. J. Winton, O. H. Yilmaz, C. S. O. Attolini, I. Gut, E. Batlle, Mex3a Marks a Slowly Dividing Subpopulation of Lgr5+ Intestinal Stem Cells. *Cell Stem Cell* **20** (2017), doi:10.1016/j.stem.2017.02.007.

12. C. S. Smillie, M. Biton, J. Ordovas-Montanes, K. M. Sullivan, G. Burgin, D. B. Graham, R. H. Herbst, N. Rogel, M. Slyper, J. Waldman, M. Sud, E. Andrews, G. Velonias, A. L. Haber, K. Jagadeesh, S. Vickovic, J. Yao, C. Stevens, D. Dionne, L. T. Nguyen, A.-C. Villani, M. Hofree, E. A. Creasey, H. Huang, O. Rozenblatt-Rosen, J. J. Garber, H. Khalili, A. N. Desch, M. J. Daly, A. N. Ananthakrishnan, A. K. Shalek, R. J. Xavier, A. Regev, Intra- and Inter-cellular Rewiring of the Human Colon during Ulcerative Colitis. *Cell* **178**, 714-730.e22 (2019).

13. J. Muñoz, D. E. Stange, A. G. Schepers, M. van de Wetering, B.-K. Koo, S. Itzkovitz, R. Volckmann, K. S. Kung, J. Koster, S. Radulescu, K. Myant, R. Versteeg, O. J. Sansom, J. H. van Es, N. Barker, A. van Oudenaarden, S. Mohammed, A. J. R. Heck, H. Clevers, The Lgr5 intestinal stem cell signature: robust expression of proposed quiescent "+4" cell markers. *EMBO J* **31**, 3079–3091 (2012).

14. I. Joanito, P. Wirapati, N. Zhao, Z. Nawaz, G. Yeo, F. Lee, C. L. P. Eng, D. C. Macalinao, M. Kahraman, H. Srinivasan, V. Lakshmanan, S. Verbandt, P. Tsantoulis, N. Gunn, P. N. Venkatesh, Z. W. Poh, R. Nahar, H. L. J. Oh, J. M. Loo, S. Chia, L. F. Cheow, E. Cheruba, M. T. Wong, L. Kua, C. Chua, A. Nguyen, J. Golovan, A. Gan, W. J. Lim, Y. A. Guo, C. K. Yap, B. Tay, Y. Hong, D. Q. Chong, A. Y. Chok, W. Y. Park, S. Han, M. H. Chang, I. Seow-En, C. Fu, R. Mathew, E. L. Toh, L. Z. Hong, A. J. Skanderup, R. DasGupta, C. A. J. Ong, K. H. Lim, E. K. W. Tan, S. L. Koo, W. Q. Leow, S. Tejpar, S. Prabhakar, I. B. Tan, Single-cell and bulk transcriptome sequencing identifies two epithelial tumor cell states and refines the consensus molecular classification of colorectal cancer. *Nature Genetics 2022 54:7* **54**, 963–975 (2022).

15. C. Cortina, G. Turon, D. Stork, X. Hernando-Momblona, M. Sevillano, M. Aguilera, S. Tosi, A. Merlos-Suárez, C. Stephan-Otto Attolini, E. Sancho, E. Batlle, A genome editing approach to study cancer stem cells in human tumors. *EMBO Mol Med* **9** (2017), doi:10.15252/emmm.201707550.

16. Z. Qi, Y. Li, B. Zhao, C. Xu, Y. Liu, H. Li, B. Zhang, X. Wang, X. Yang, W. Xie, B. Li, J. D. J. Han, Y. G. Chen, BMP restricts stemness of intestinal Lgr5 + stem cells by directly suppressing their signature genes. *Nat Commun* **8** (2017), doi:10.1038/ncomms13824.

17. A. Ayyaz, S. Kumar, B. Sangiorgi, B. Ghoshal, J. Gosio, S. Ouladan, M. Fink, S. Barutcu, D. Trcka, J. Shen, K. Chan, J. L. Wrana, A. Gregorieff, Single-cell transcriptomes of the regenerating intestine reveal a revival stem cell. *Nature* **569** (2019), doi:10.1038/s41586-019-1154-y.
